# Behavioural relevance of spontaneous, transient brain network interactions in fMRI

**DOI:** 10.1101/779736

**Authors:** D. Vidaurre, A. Llera, S.M. Smith, M.W. Woolrich

**Affiliations:** University of Oxford; Donders Institute

## Abstract

How spontaneously fluctuating functional magnetic resonance imaging (fMRI) signals in different brain regions relate to behaviour has been an open question for decades. Correlations in these signals, known as functional connectivity, can be averaged over several minutes of data to provide a stable representation of the functional network architecture for an individual. However, associations between these stable features and behavioural traits have been shown to be dominated by individual differences in anatomy. Here, using kernel learning tools, we propose methods to assess and compare the relation between time-varying functional connectivity, time-averaged functional connectivity, structural brain data, and non-imaging subject behavioural traits. We applied these methods on Human Connectome Project resting-state fMRI data to show that time-varying fMRI functional connectivity, detected at time-scales of a few seconds, has associations with some behavioural traits that are not dominated by anatomy. Despite time-averaged functional connectivity accounting for the largest proportion of variability in the fMRI signal between individuals, we found that some aspects of intelligence could only be explained by time-varying functional connectivity. The finding that time-varying fMRI functional connectivity has a unique relationship to population behavioural variability suggests that it might reflect transient neuronal communication fluctuating around a stable neural architecture.

**Significance statement:** Complex cognition is dynamic and emerges from the interaction between multiple areas across the whole brain, i.e. from brain networks. Hence, the utility of functional MRI to investigate brain activity depends on how well it can capture time-varying network interactions. Here, we develop methods to predict behavioural traits of individuals from either time-varying functional connectivity, time-averaged functional connectivity, or structural brain data. We use these to show that the time-varying nature of functional brain networks in fMRI can be reliably measured and can explain aspects of behaviour not captured by structural data or time-averaged functional connectivity. These results provide important insights to the question of how the brain represents information and how these representations can be measured with fMRI.

## Introduction

The emergence of large-scale distributed networks in spontaneous brain activity as measured by functional magnetic resonance imaging (fMRI) is a widely-studied phenomenon (Biswal et al., 1995; Fox and Raichle, 2007). These networks have been consistently identified using cross-regional temporal correlations – referred to as functional connectivity (FC) (Damoiseaux et al., 2006; Smith et al., 2013; Hipp and Siegel, 2015). Typically, FC is estimated by averaging over several minutes of data (e.g. across a scanning session, for each pair of regions) to provide a stable representation of the functional network architecture for an individual (Finn et al., 2015). This *time-averaged FC* has previously been associated with mental performance (Hampson et al., 2006; Hasson et al., 2009) and, more generally, to widespread behavioural phenotypes (Smith et al., 2015). However, there is evidence that some of these associations are to a large extent driven by structural differences between subjects (Bijsterbosch et al., 2018; Llera et al., 2019). We hypothesised that, while time-averaged FC might to some extent be dominated by structural information, the temporal deviations of FC might be less so, and could thereby have a distinct relationship with behaviour. This would provide evidence that time-varying FC from fMRI can reflect momentary neuronal communication fluctuating around a stable functional architecture, and might be related to dynamic elements of cognition such as attention and thinking (Smallwood and Schooler, 2015; Kucyi, 2017).

While there is clear evidence that electrophysiologically-derived FC relates to momentary mental states (Palva and Palva, 2012; Hipp et al., 2011; O’Neill et al., 2017; Quinn et al., 2018), whether dynamic changes in fMRI-derived FC reflect distinct and transient patterns of communication between neuronal populations is still controversial (Gratton et al., 2018). In the absence of stimuli, measures of ongoing behavioural outputs or any ground truth, discerning whether time-varying FC carries biological meaning in the resting state is indeed not straightforward (Lurie et al., 2018; Kucyi et al., 2018). One possibility is to use indirect behavioural correlates, for example, by assessing the extent to which FC prior to task onset influences task performance (Sadaghiani et al., 2015), quantifying how the execution of a task induces differences in subsequent resting-state FC (Waites et al., 2005), or using a low demanding task with well-defined behavioural information as a surrogate of actual resting-state (Kucyi et al., 2017). However, these are normally subtle effects, and other researchers have reported little or no differences in FC between task and rest (Hampson et al., 2006; Gratton et al., 2018).

Here, we take a different route, by relating time-varying FC to population variability in behavioural traits. For this purpose, we implemented a framework to predict subject behavioural traits from either time-varying FC, time-averaged FC, or structural data. Critically, this was done in such a way that the prediction could be abstracted from the very distinct nature of the features used to represent each of the three modalities, allowing us to compare their relative and unique contribution to the prediction in an unbiased manner. Using different groups of behavioural traits, we used this approach to explore the relationship between population behaviour, time-averaged FC and time-varying FC, after accounting for the explanatory power of the structural connectivity features. We reasoned that if fMRI time-varying FC represents biologically meaningful communication between neuronal populations, then it should be capable of accounting for aspects of the subjects’ behavioural phenotypes not explained by the time-averaged FC or the structural information. We found that this was the case, particularly for the traits that are generally related to intelligence.

To measure time-varying fMRI FC, we used a state-based model where each state is associated with a specific pattern of FC (Vidaurre et al., 2017), such that instantaneous changes in FC manifest as a change of state. This approach is based on a version of the hidden Markov model (HMM) that, in comparison to previous versions of the HMM used on fMRI (Vidaurre et al., 2017; Stevner et al., 2019; Baldassano et al., 2017; Shappell et al., 2019; Zhang et al., 2019), emphasises changes in FC over changes in amplitude. To model each subject, the HMM uses a temporally-organised mixture of (quasi-) stable FC patterns in the form of region-by-region covariance matrices. This is an appropriate choice to compare time-varying FC with time-averaged FC, since time-averaged FC is also based on region-by-region covariance matrices. To model structural variability, we used fractional anisotropy (FA; Basser and Pierpaoli, 1996), mean diffusivity (MD; Basser et al., 1994) and voxel-based morphometry (VBM; Ashburner and Friston, 2000).

## Results

In this section, we first summarise the basic steps of the analysis, which are presented in more detail in Methods, and then go on to show that there are aspects of behaviour that are uniquely expressed in both time-averaged and time-varying FC. We also show how each of these representations explicitly relate to each other, and to the structural data, in terms of their relation to behaviour. Overall, these analyses suggest that time-averaged and time-varying FC can indeed reflect separate aspects of brain activity.

### Functional representations of the data

We used 1003 subjects’ resting-state fMRI data with TR=750ms from the Human Connectome Project (HCP; Smith et al, 2013b), where each subject underwent four 15-min sessions (two per day). We used a data-driven parcellation obtained through spatial independent component analysis (ICA), and extracted 50 components (Beckmann et al., 2009). The time series of these ICA components were then standardised separately for each session.

We considered two different FC-related representations of the data. The first representation is a time-averaged FC model, where we represented each subject as one (50 by 50) Pearson’s correlation matrix across all ICA component time series (Smith et al, 2013). Because the time series have unit-variance, these correlation matrices are equivalent to the corresponding covariance matrices.

The second representation corresponds to a time-varying FC model, where the ICA time series were fed to a hidden Markov model (HMM), which we first ran at the group level – i.e. on the concatenated time series for all subjects. The HMM represents the data as (i) a collection of states, each represented by a certain probability distribution; (ii) time series of state activation probabilities, one per state and time point, referred to as state time courses; and (iii) a transition probability matrix containing the probability of switching from one state to another within a session (Vidaurre et al, 2017; Vidaurre et al, 2018a); see Methods and **Fig SI-1** for an illustration. As opposed e.g. to Vidaurre et al. 2017 or Baker et al. 2014, which represented states using ordinary Gaussian distributions, here we implemented an HMM designed to emphasise periods in time with distinct FC (also see **Fig SI-2** for a graphical, exemplary comparison). Specifically, each HMM state is represented by a covariance matrix across ICA components, so changes of state activations within session –expressed by the state time courses– correspond to modulations of covariance above and beyond the average covariance or FC. In this model, the state-specific covariances and transition probability matrix are estimated at the group level, whereas the state time courses are subject-specific (Vidaurre et al, 2017). Akin to the procedure known as dual regression in ICA (Nickerson et al., 2017), we then performed a process of *dual-estimation* to obtain subject-specific versions of the group-level HMM in order to get a fuller subject-specific description of time-varying FC, where each subject has their own set of state-specific covariances (i.e., FC matrices), transition probability matrix, and state time courses.

We trained the models with eight states, without optimising for this number; previous work, however, suggested that the relations to behaviour are relatively robust across a reasonable range of states (Vidaurre et al., 2017). As often occurs with other models where the estimation depends on an optimisation process, the inference of the HMM can potentially produce different solutions depending on the initialisation (Vidaurre et al, 2019). Thus, in order to ensure that our conclusions were robust, we conducted five repetitions of the inference.

### Prediction of behavioural variability

From the two functional representations just described, and the three considered anatomical descriptors (FA, MD, and VBM), we went on to assess how each of these can predict the considered behavioural traits. Within a 10-fold cross-validation scheme that respected the family structure of the HCP data (Winkler et al., 2015) by never splitting families between folds, we predicted a number of behavioural traits within six different groups of variables: demographic, cognitive, affective, personality and sleep-related (**Table SI-1**). The word “behavioural” is used here in a general sense, even though we included demographic and life-factor traits that are not purely behavioural.

For predicting behaviour, we used an approach based on distance matrices (DM) and cross-validated, motion-corrected kernel ridge regression (KRR; Saunders et al, 1999; Schölkopf and Smola, 2001; He et al., 2020). Specifically, we computed (*N* by *N*) distance matrices (DM), where *N* is the number of subjects and where the distances are calculated to capture how different a specific representation is between each pair of subjects. Overall, there is one representation for the time-averaged FC (i.e. the FC time-averaged network matrix), five representations for the HMM-based FC (i.e. one per repetition of the HMM inference), and one representation for each of the structural information measures (FA, MD, VBM); this yielded one time-averaged-FC-DM, five HMM-DMs (one per repetition of the inference), and three structural-DMs (FA-DM, MD-DM and VBM-DM). Therefore, whereas the approach to compute the distances is specific to each modality, all modalities end up being reduced to the same format (a DM); see Methods for details about how the pairwise distances for each modality were computed. The motivation of the KRR approach is two-fold. First, because the prediction is based exclusively on distances, we can decide on a sensible distance measure to use between different representations, instead of manually deciding what features to use to represent a representation. This offers a clean solution to the problem of how to make predictions using a complex object like an HMM, which it is not obvious how to convert into a vector of representative features; instead, using an appropriate measure to quantify how different two HMMs are is relatively straightforward. Second, having all types of representation (time-averaged FC, HMM or structural) in the same format (a DM) makes it easier to compare the explanatory power of each modality in predicting the subject traits, which otherwise could be heavily dependent on their specific parameterisation. See Methods for a mathematical description of KRR.

With the goal of exploring the influence of the structural information on the functional representations, we ran the predictions on the uncorrected behavioural traits, as well as on the behavioural traits after regressing out (deconfounding) the structural (FA, DM or VBM) information; see Methods for details. A scheme of the prediction procedure is illustrated in **Fig 1** for the dual-estimated HMMs: on top, the prediction from the structural information; at the bottom, the subsequent structure-deconfounded estimation from the dual-estimated HMM. The estimation for the time-averaged FC is analogous.

**Fig 1.**
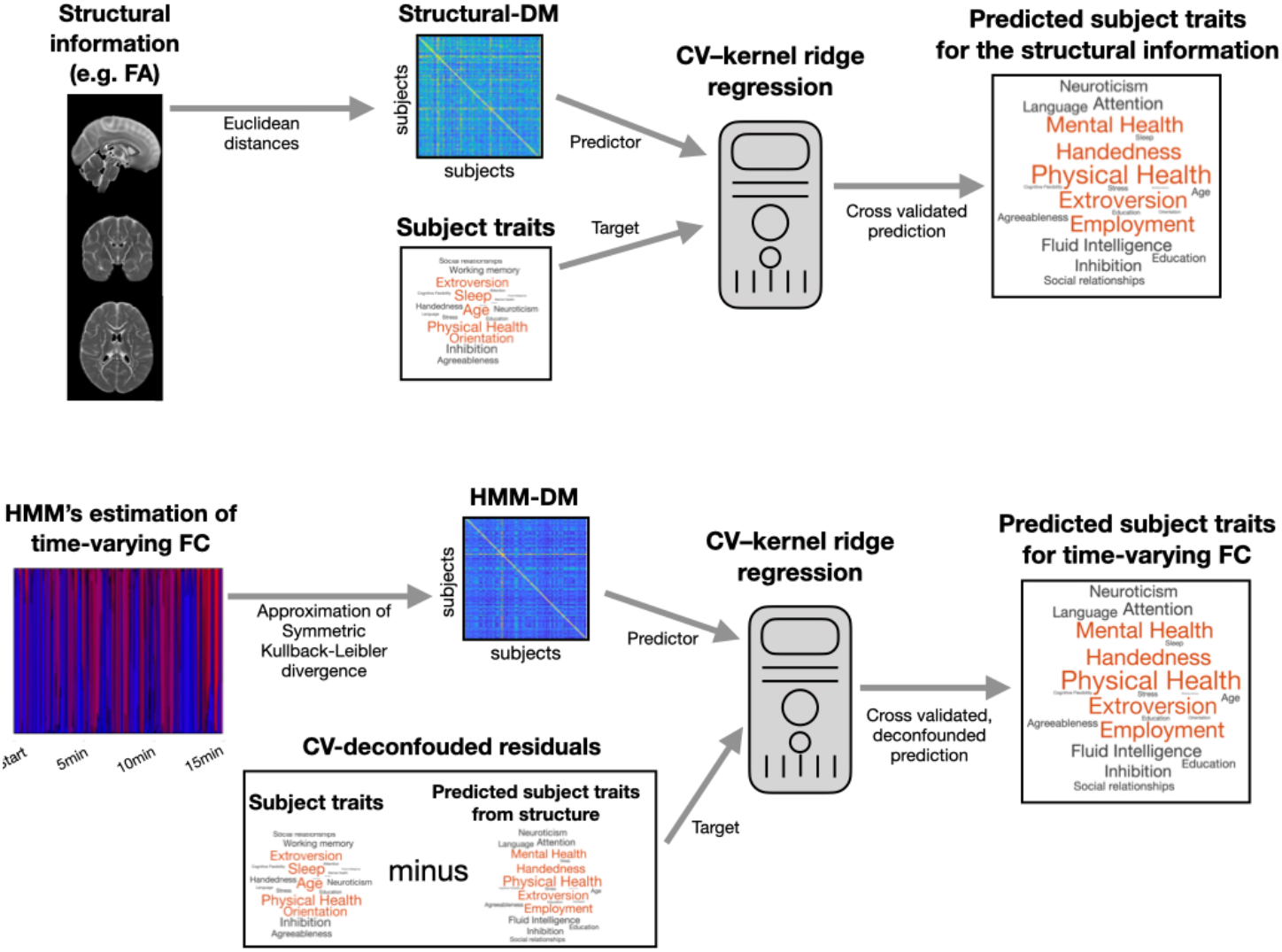
Prediction scheme using representations in terms of distance matrices (DM). On top, cross-validated prediction from the structural information; at the bottom, structure-deconfounded prediction from the dual-estimated HMM, which contains information of time-varying FC. The CV-deconfounded residuals are the traits after discounting the influence of the structural information. An analogous procedure is used for the time-averaged FC.

### Time-varying FC explains distinct aspects of behaviour

Taking into account the structural information, we next show that time-varying FC contains information from some behavioural traits that is not contained in the time-averaged FC, and vice versa that the time-averaged FC is a better predictor than the time-varying FC representation for other behavioural traits.

**Fig 2** shows a comparison of the prediction performances between the HMM representation and the time-averaged FC representation for the six behavioural groups listed in **Table SI-1**. This is presented for both the structure-deconfounded (i.e. for FA, MD and VBM; see above for details about deconfounding) and the non-deconfounded case. The top panels present the cross-validated explained variance (*r*^*2*^, computed as a squared Pearson’s correlation) for the HMM and time-averaged FC representation; statistical significance through Bonferroni-corrected parametric testing is indicated by colour. Note that although the predictions are not very high (Kong et al., 2019; Pervaiz et al, 2020), several are still significant. The middle panels reflect the difference between the two – which is positive when the HMM representation is a better predictor and negative otherwise; and the bottom panels contain an average of these differences per behavioural group. Statistical significance of whether one representation has a higher *r*^*2*^ than the other across traits is indicated within the bottom panels (*<0.05; **<0.01; permutation testing) for each behavioural group. Note that, as mentioned above, in the case of the HMM there are five different predictions per trait, one per run of the HMM inference; therefore, also, there are five prediction differences between the HMM and the time-averaged representation per trait, and five dots per trait in the middle panels. As observed, there is considerable variability in which type of representation (HMM- or time-averaged-FC-based) represents the traits better. Also, structure-deconfounding affects the prediction accuracy considerably, confirming previous studies on the influence of the structural information on FC-based predictions (Bijsterbosch et al., 2018; Llera et al., 2019). In this regard, **Fig SI-3** shows the loss of accuracy in percentage after correcting for the structure for each modality, grouped by behavioural group. For reference, **Fig SI-4** shows the (uncorrected) explained variance by each structural representation.

**Fig 2.**
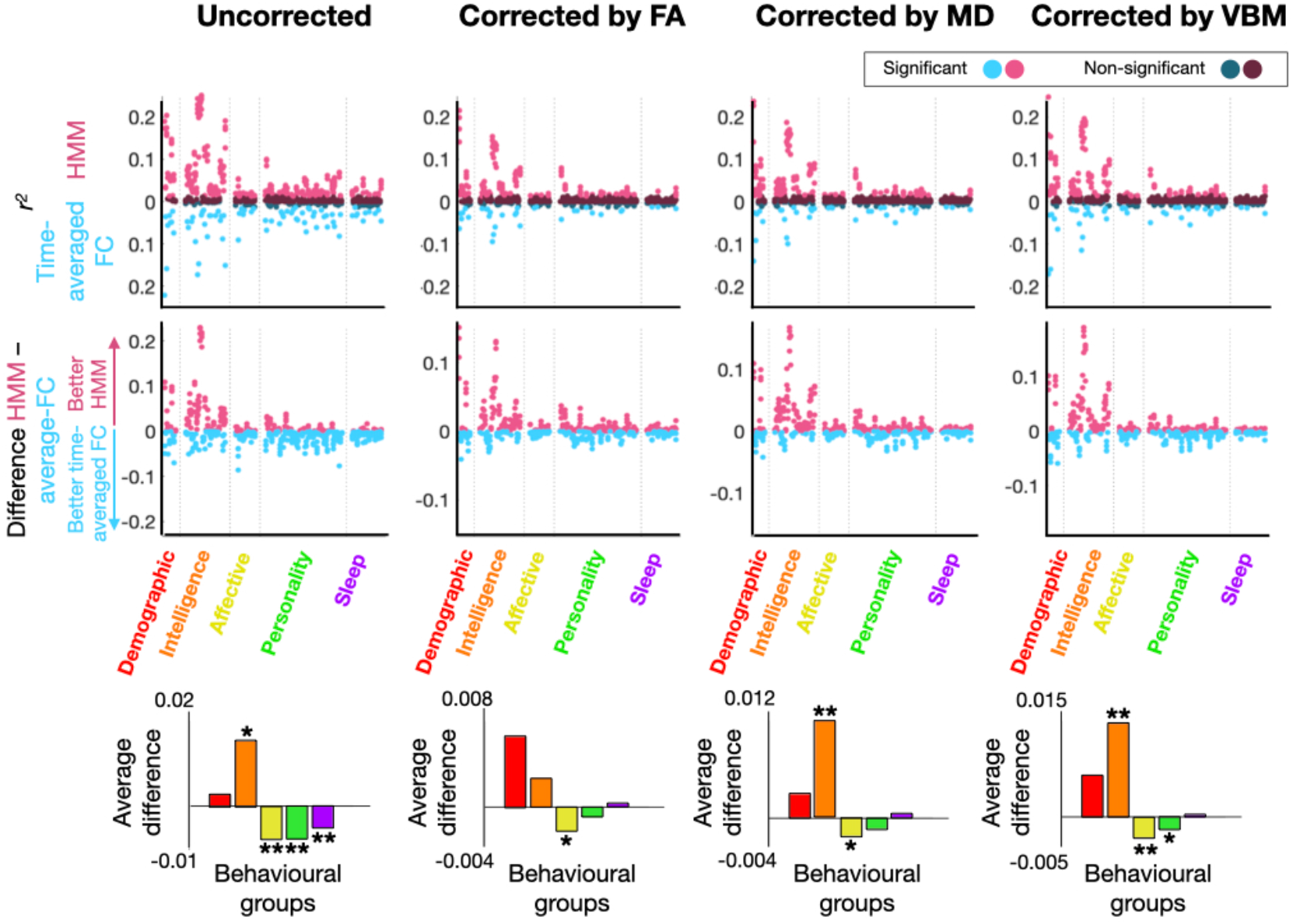
Explained variance *r*^*2*^ (in terms of squared Pearson’s correlation) for the prediction of behavioural traits using the time-averaged-FC-DM and the HMM-DMs. In the top panels, *r*^*2*^ values (upwards for the HMM and downwards for the time-averaged representation; lighter colours represent statistically significant predictions and darker non-significant; Bonferroni-corrected parametric testing); in the middle panels, difference between the HMM and the time-average FC representations; in the bottom panels, the average differences aggregated by behavioural group (* and ** reflect statistical significance for significance levels of 0.05 and 0.01; permutation testing).

From this analysis, two conclusions are apparent. First and most importantly, that the behavioural groups are well separated by which representation is more effective in predicting them, with intelligence being particularly well predicted by the HMM representations. Second, that correcting by the structural information improves the relative performance of the HMM-DM compared to the time-averaged-FC-DM (see also **Fig SI-3**).

### Changes in variance and amplitude of the signal do not explain behaviour

In order to investigate the possibility that the predictions are primarily driven by within-session changes in the variance or amplitude of the signal instead of FC, we ran two additional varieties of the HMM. These will be compared with the FC-based version of the HMM used throughout the paper –here referred to as FC-HMM–, where each state is parametrised as a Gaussian distribution with zero mean and a full covariance matrix. In the first of the new varieties, referred to as mean-HMM, the states where characterised by Gaussian distributions with distinct patterns of signal amplitude (encoded in the mean parameter), and a common full covariance matrix shared across states. In the second, the var-HMM, the states were characterised by Gaussian distributions with a diagonal covariance matrix and zero mean, modelling just variance and not actual covariance between regions. In these models, the FC was not allowed to vary between states. Furthermore, while the mean-HMM takes into consideration the time-averaged FC through the shared covariance matrix, the var-HMM does not model FC at all. **Fig 3** presents the explained variance of the FC-HMM versus the explained variance of each of the other two HMM varieties. As observed, the explained variance for FC-HMM is consistently superior, highlighting the importance of accounting for time-varying FC above and beyond changes in amplitude and variance. Note that, although these models differ on the number of parameters and their complexity –which in principle could influence the quality of the predictions– here we are abstracting ourselves of these differences by performing the predictions on the base of DMs only. This means that the KRR models have always the same number of parameters regardless of the modality (see Methods).

**Fig 3.**
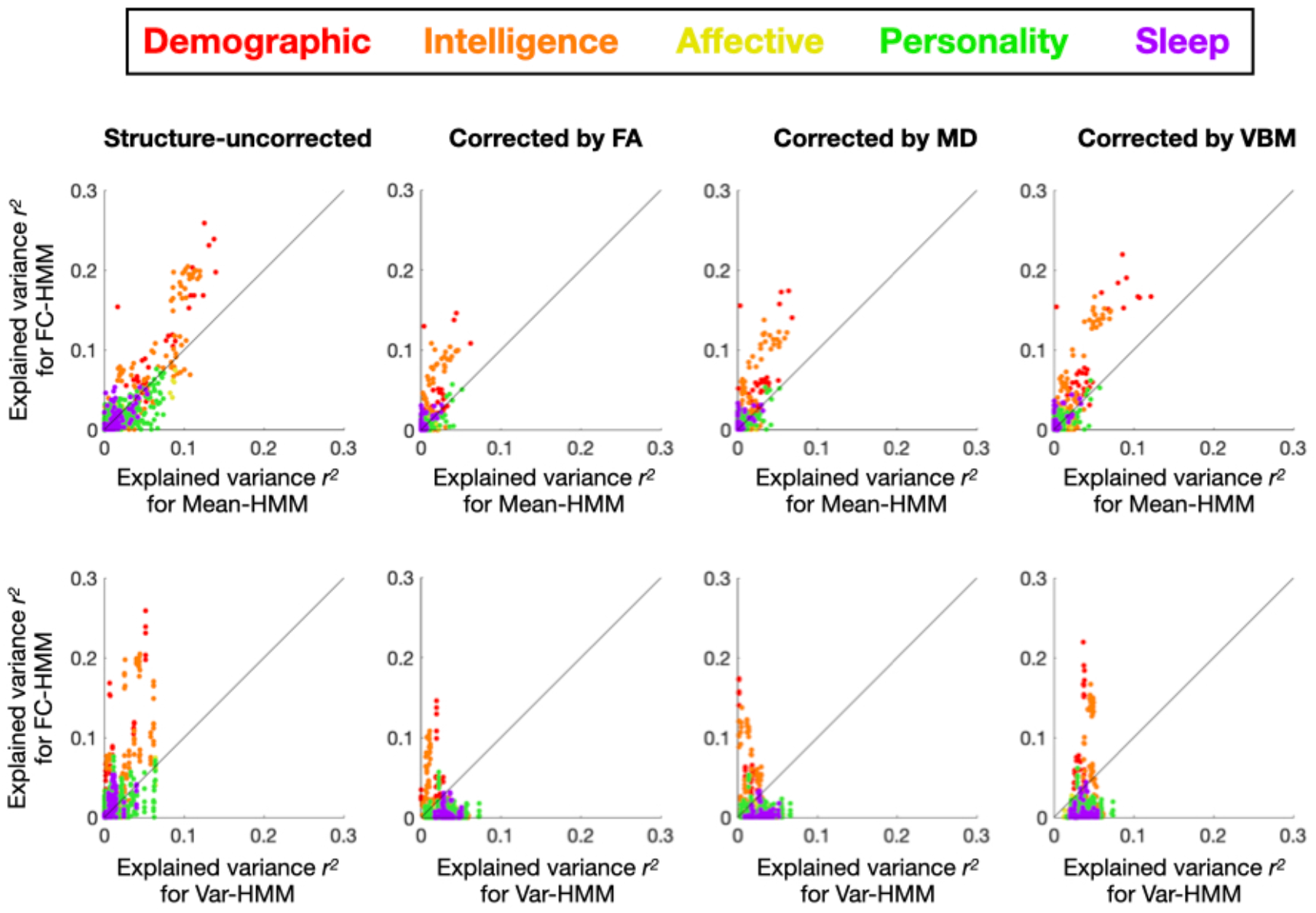
Behavioural explained variance *r*^*2*^ (defined as squared Pearson’s correlation) by the FC-based HMM model (which is the type of HMM used throughout the paper, Y-axis) vs (i) the mean-HMM, a type of HMM with one shared covariance matrix and one “mean” parameter per state that models changes in amplitude (top row; X-axis), and (ii) the Var-HMM, a type of HMM with state-specific variance parameters, i.e. with no cross-region covariances (bottom row; X-axis).

### Time-varying FC is more dissimilar to the structural information than time-averaged FC

Through their differences in prediction accuracy of traits, we have investigated the amount of information contained in either the time-averaged or the time-varying FC representations with respect to behaviour. A complementary question is to what extent, specifically, do these brain representations contain similar or different information about behaviour. That is, if two representations are very similar with respect to a given behavioural group, that means that they represent similar information about that specific aspect of behaviour; if they are very dissimilar, it means that their information about the behavioural group is mostly non-overlapping.

For each group of behavioural traits (see **Table SI-1**), we correlated the trait predictions between each pair of brain representations: time-averaged FC, each of the three structural measures, and each of the five HMM runs for the three HMM configurations described in the previous section (i.e. FC-HMM, which is the main model used throughout this study; mean-HMM, which only models the amplitude; and var-HMM, which models changes in variance). In the spirit of the *Representation Similarity Analysis* literature (Kriegeskorte et al., 2008), this procedure produced a (no. of brain representations by no. of brain representations) similarity matrix per behavioural group, capturing how correlated the prediction of the behavioural traits was between each pair of representations.

**Fig 4A** presents the corresponding similarity matrices, for each behavioural group. The five matrices have some common patterns but some differences are also apparent. The most relevant pattern here is that, in all cases, the structural representations were much more related (in terms of explaining behaviour) to the time-averaged FC than to the FC-HMM representations, confirming that time-varying FC is more unrelated to the structure than time-averaged FC. This is further explored in **Fig 4B**, where we show the probability density of the corresponding correlations in terms of explaining behaviour between the structural representations on the one hand and either the FC-HMM (blue) or the time-averaged FC (red) representations on the other hand. The probability densities of the correlations were estimated by bootstrapping (Efron and Tibshirani, 1986). As observed, the differences are large and significant. We can also observe that the five FC-HMM representations are more consistent in explaining the demographic traits (i.e. the correlation between HMM runs is higher) than they are in the other behavioural groups, and they are also more related to the time-averaged FC representation for this behavioural group than for the others. The latter point can be seen in **Fig 4C**, which shows the probability density of correlations (estimated by bootstrapping) between the FC-HMM and the time-averaged FC in terms of how they explain each behavioural group. The larger similarity for the demographic group is apparent. The other two types of HMM estimations, having fewer parameters and without capturing any information about time-varying FC, are in general more similar across runs and quite different from the FC-HMM, indicating that the FC-HMM approach is unlikely to be purely driven by changes in amplitude. In terms of the structural information, MD and FA are fairly similar to each other for all behavioural groups, but their similarity to VBM varies according to the behavioural group (highest for intelligence and sleep; lowest for demography).

**Fig 4.**
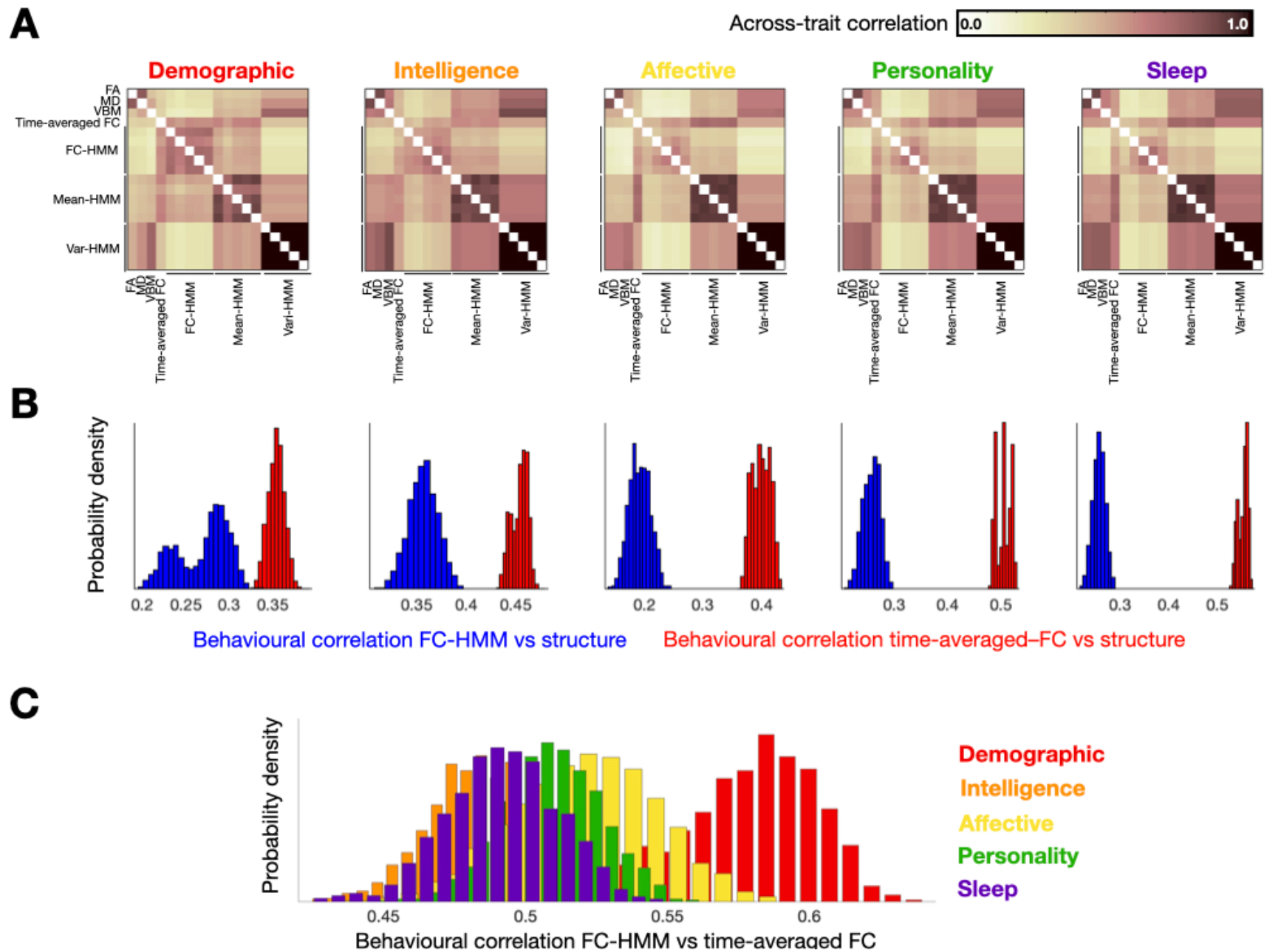
**A**. How similar are the different representations in explaining behaviour? Similarity matrices (in terms of Pearson’s correlation) capturing how similar the prediction of behavioural traits was between each pair of representations are shown for each of the five behavioural groups: time-averaged FC, HMM-based representations including time-varying FC (FC-HMM; used throughout the paper), HMM representations including only changes in amplitude (mean-HMM) or variance (var-HMM), and structural (FA, MD and VBM). **B**. Distribution densities (obtained via bootstrapping) of between-modality correlations (in terms of explaining behaviour) show that the time-averaged FC representation is more related to the structural representations than the time-varying FC. **C**. The correlations between the time-varying and the time-averaged FC representations are higher for the demographic traits than for the other behavioural groups.

Altogether, this analysis revealed clear differences and similarities between the different neuroimaging representations in terms of explaining behaviour, and provides further evidence that time-varying FC is more unrelated to the anatomy than the time-averaged FC.

### Reproducibility of DMs

The reproducibility of the estimation of both the representations and the behavioural predictions can be relevant in terms, for example, of a clinical application. In the previous section, we have considered how different estimations of the various representations differ in their relation to behaviour. Here, we analyse another aspect the representations’ reproducibility: how robust are these representations, per se, across scanning sessions.

The HCP data contains four sessions per subject, with the first two (1 and 2) being acquired on one day and the last two (3 and 4) on the following day. In order to further quantify the reproducibility of the estimations, we estimated separate time-averaged FC and (FC-)HMM models for the first day and for the second day, i.e. for sessions 1 and 2, and then for sessions 3 and 4. We also estimated models for the first session of the day, and then separately for the second session of the day. For each of these two half-split estimations, (HMM- or time-averaged FC-related), we then computed DMs. **Fig 5** presents a quantitative assessment of the reproducibility of the estimations in terms of how similar their respective DMs were across half-splits of the data. Here, the dots represent a measure of distance between one pair of subjects.

**Fig 5.**
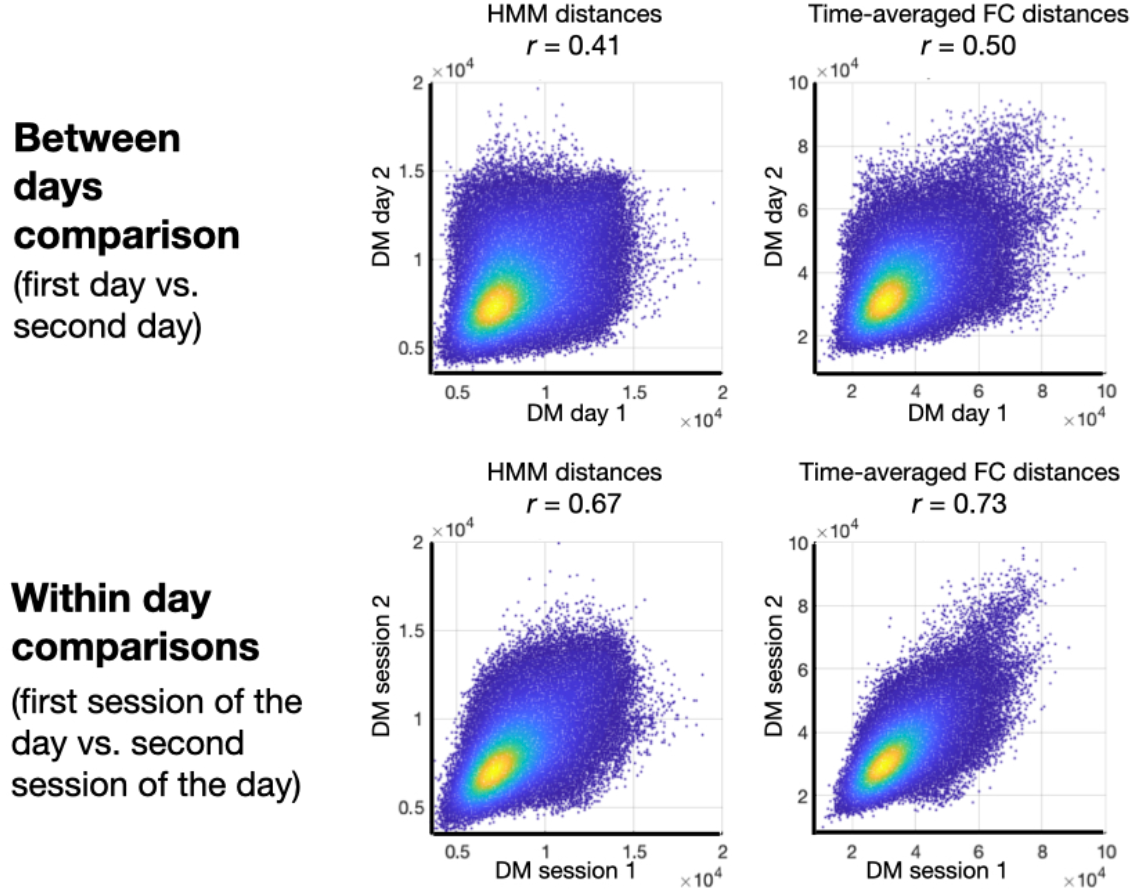
Reproducibility of the estimations between the first and the second day of scanning (top), and between the first session and the second session of each day (bottom), for the HMM (left) and the time-averaged FC representation (right). Each dot corresponds to an element of the DM, i.e. a distance measure between a pair of subjects, and the colour reflect the density of dots. For each panel, the correlation between the DMs (i.e. across dots) is reported as a Pearson’s correlation *r*.

As expected, the reproducibility within day is considerably larger than between days for both types of representation. Importantly, the time-averaged FC description (being a simpler quantity to estimate) exhibits in general a higher between-session reproducibility than the HMM representation (Vidaurre et al. 2019). This can be due to the time-averaged FC not just being a simpler quantity to estimate, but also to the HMM being potentially better able to capture session-specific information thanks to its time-resolved nature.

## Discussion

In resting state fMRI, the quantification of time-varying functional connectivity (FC) has elicited considerable interest and controversy: that is, to what extent can we measure and interpret within-session changes in the patterns of FC between areas? Whereas many studies rely on the average magnitude of activation that is evoked by a task or stimulus, FC is a second-order statistic and therefore is harder to estimate accurately. Similarly, it is unclear whether FC can reflect changing patterns of communication between distant neuronal populations, and therefore be meaningful for investigating cognition. Even though the total amount of between-subject variability attributed to stable subject FC features (i.e. that do not change within session and are preserved for each subject across sessions) is considerably higher than the within-session variability (i.e. that change within a session; Gratton et al., 2018), here we show that fMRI-derived FC indeed contains both stable and time-varying behaviourally meaningful information, and that time-varying FC can explain behavioural variability that is less likely to be mediated by structural connectivity and other anatomical features. This suggests that time-varying FC may represent meaningful neuronal dynamics related to certain aspects of behaviour. As a consequence, the study of FC fluctuations remains promising for the understanding of transient cognition.

To answer this question, it is informative to disentangle the different mechanisms by which time-varying FC computed from fMRI data could be non-informative: first, the characterisation of time-varying FC may be limited by fundamentally technical issues; second, the actual amount of information contained in the time variations, when assessed unbiasedly, may be negligible; and third, even if we can prove that there is non-negligible information in time-varying FC that can be reliably quantified, it may still not be cognitively significant. We argue that certain technical limitations do not apply to all methods of estimating time-varying FC equally. In the case of the HMM, for example, the technical limitation of having a statistically unstable estimation due to limited amounts of data (e.g. when using sliding windows) is overcome by using large amounts of data in the estimation of each state through the ability to pool over all the data from repeated visits to the state (on average, 125h per state in the present data set).

It has been shown that the time-averaged (subject-specific) FC features represent most of the variance in fMRI data (Gratton et al., 2018). However, the fact that time-varying FC explains considerably less variance does not necessarily mean that time-varying FC is deficient in explaining behavioural traits. We consider that discussing the physiological relevance of a brain representation in terms of explained variance (of the data) is not appropriate for two reasons: (i) that “physiological relevance” must be connected to a specific scientific question – e.g. relevant to the study of attention; and (ii) that, provided such a question, there is not prior evidence that the most informative aspect of the signal (for that question) is the one that explains the most variance in the data. For example, in the context of prediction it is a well-known phenomenon that the first principal components of the predictor data are not necessarily the most explanatory to predict the target variable (Frank and Friedmann, 1993). As an example closer to neuroscience, electrophysiological signals hold most of their variance at lower frequencies. In comparison, only a small fraction of variance is contained e.g. in the gamma frequencies (>40 Hz). These, however, have been demonstrated to be essential to behaviour (Jensen et al., 2007). In summary, the argument that there is considerably more variability in the between-subject than in the within-subject differences cannot be used to claim the lack of biological relevance of these features.

### Relation to previous work

Some of the conclusions of this study relate to the recent work from Liégeois et al. (2019), who found, in fMRI, that the autoregressive model (which linearly represents how on average the signal at time point *t* depends across regions on the signal at time point *t*-1) was often more explanatory of behavioural variability than the standard time-averaged FC estimation. Because the autoregressive model is known to describe the dynamics of the signal well (Liégeois et al., 2017), the conclusion of this study was that the dynamic aspects of the data can often explain behaviour better than (average) instantaneous fMRI correlations. Critically, there is a conceptual distinction between a model of the multivariate dynamics of the system (as captured by the autoregressive model) and time-varying FC (as captured by the HMM) that is important to the message of this study. Specifically, while both the HMM and the autoregressive model can capture time-varying FC, the autoregressive model also captures other elements such as spectral information (Vidaurre et al., 2016), while the HMM captures aspects of the data that the autoregressive model does not explicitly account for, such as the identification of time-resolved transient events. Therefore, the autoregressive model is not able to answer our question, which is focussed specifically on FC: i.e., do variations over time in the fMRI FC have biological significance above and beyond the temporally averaged FC, and also the structural information? This question is important as it speaks to the extent to which FC can represent instantaneous neural communication. These questions require a model that explicitly considers variations around the time-averaged FC in a way that is not mixed with these other elements. The version of the HMM used here is one way to achieve this, but not the only one. Other data descriptions capturing related or different aspects could also be considered, such as those based on signal events (Allan et al., 2014) or quasi-periodic patterns (Thompson et al., 2014).

It is worth noticing that, in accordance with the growing body of work on predicting behavioural traits from functional connectivity on the HCP data, the predictions were modest; see Smith et al. (2016), Kong et al. (2019), Greene et al. (2018) and Pervaiz et al. (2020) among many others. However, these are still clearly significant (Smith et al, 2014; Vidaurre at al., 2017), allowing us to disentangle the time-varying from the time-averaged FC behavioural relevance in terms of trait prediction. Future work will aim at replicating these results on the UK Biobank, where higher prediction accuracies have been observed (Pervaiz et al., 2020).

### Limitations and open questions

It is also important to appreciate that neither the HMM nor other commonly used time-varying FC estimators are explicitly biophysical models. Decisions about the appropriate number of states and other parameters are useful insofar as they affect the extent to which we can address the specific question at hand. For example, estimating more states will offer a more fine-grained representation of the data, which might be necessary in certain applications but cannot be interpreted as more or less faithful to the biology. In general, different parametrisations just offer different perspectives of the data, and, assuming model identifiability, the HMM is not more or less valid than other models. We also acknowledge that, while the version of the HMM used in this work is designed to emphasise time-varying FC, it could also be sensitive to changes in amplitude (Duff et al., 2018). However, we have explicitly tested that a version of the HMM only based on changes in amplitude is unable to explain behaviour to the same extent, emphasising the importance of time-varying FC. Other aspects of the data that can influence the HMM estimation are long-range temporal dependencies, which are not explicitly modelled by the HMM (Shappell et al., 2019). A quantitative assessment of the long-term dependencies in the data and how they affect the HMM estimation will be subject of future work.

An important methodological consideration is that, even though all the representations are unbiasedly compared at the level of prediction because of their common DM representation, our analysis still depends on the choice of how to compute the distances. For example, in this study we used a Kullback-Leibler divergence approximation to compute distances between HMM representations (Do, 2003; see Methods). Alternatively, one could compute differences purely based on the temporal aspects of the model (e.g. the transition probability matrix) or its spatial properties. Related to this point, the merits of kernel-based approaches come at the expense of neuroanatomical interpretation: since we no longer have one regression coefficient per spatial area, but one regression coefficient per subject (see Methods), and given also that the distances matrices (on which the prediction is based) are computed in an unsupervised fashion, it is not straightforward to see which areas have actually driven the prediction. Future work will address these questions, including how to compute between-model distances as part of the prediction and in an interpretable manner, so that the most predictive features of the models are identified in a data-driven way.

One aspect to consider about models for which inversion does not have a mathematically closed formulation (as is the case of ICA and the HMM among many others – but not of Pearson’s correlation or the autoregressive model) is the fact that, every time we estimate the model, we might get a slightly different description of the data insofar as the estimation has a random seed (see for example **Fig 4**). Even though the HMM inference is relatively stable on this data set (Vidaurre et al, 2017), that is not guaranteed to be the case always. Again, these are not biophysical models, so all estimations are theoretically valid as far as they are considered as what they are: data-driven descriptions. Although there exist statistical testing approaches available to combine across estimations so that statistical power is boosted (Vidaurre et al., 2019), in this work we have analysed each estimation separately to ensure the comparability of the results.

A further caveat is that the ICA maps are known to contain important subject-specific differences that can be relevant to behaviour (Bijsterbosch et al. 2018). These differences were not considered in this paper, as we estimated both time-varying and time-averaged FC using only the ICA time series. In future work, we will study the combination of these analyses with techniques that are more suitable to account for this information (Harrison et al., 2015).

Finally, it is worth noting that, to be conservative, we have performed the (group-level) HMM estimation within the cross-validation loop. However, since the HMM estimation is completely unsupervised and does not make any use of the labels, it would have been also correct to obtain the (dual-estimated) HMMs prior to, and out of, the prediction cross-validation loop. Whether or not this is acceptable to do depends on the practicalities of the application. For example, if we wish to predict whether a new subject is going to develop a disease in the future given their brain data, it would not be a problem to rerun, on the entire data set (i.e. including the new subject), a unsupervised dimensionality reduction algorithm (like the HMM) before doing the prediction as far as such algorithm is unsupervised. Doing this would not make a diagnosis any less valid – only perhaps slower. But sometimes we would not have access to the original data at the time of prediction, in which case a proper validation of the method would need to cross-validate the HMM estimation as we did here.

In summary, this study presents methods to use different sources of brain data and/or models for prediction, in a way that makes comparisons possible in terms of their explanatory power of behavioural or clinical variables. Using this method, we have shown that time-averaged and time-varying FC explain distinct aspects of behaviour, above and beyond the behavioural variability expressed on the considered structural brain data.

## Methods

We provide some details on preprocessing, the nature of the hidden Markov model estimation and its different varieties, the computation of the distances between each pair of subjects for each of the considered measures or subject variables, and the use of kernel regression to test the relation between imaging and non-imaging variables, how we accounted for the structural information and the influence of motion.

### Preprocessing

We used the “minimal preprocessed” data from the HCP. Since the preprocessing of this data has already been described in detail (Smith et al., 2013b), we will discuss it only briefly. Structured artefact removal using independent component analysis (ICA) and FIX (Griffanti et al., 2014) removed more than 99% of the artefactual ICA components in the data set. No low-pass temporal filter was used, and only minimal high-pass filtering was applied (cutoff= 2000s), essentially removing the linear trends of the data. Since ICA-based methods have been shown to better characterise the signal than other data-driven approaches such as k-means (in particular on the HCP data; Bzdok et al., 2016), we used group spatial-ICA to obtain a “parcellation” of 50 components that covers both the cortical surfaces and the subcortical areas (without using global signal regression). This parcellation was then used to project the Fmri data into 50-dimensional time series. These time series were finally standardised separately for each scan, subject and ICA component.

### An FC-focused hidden Markov model

The hidden Markov model (HMM) is a probabilistic framework used to model time series using a finite number of recurring patterns that succeed each other in some order (Rabiner, 1989). The key assumptions of this approach are: that the data can be reasonably represented using a discrete number of probabilistic models; that occurrence of these models is exclusive – i.e. the state time courses’ summation across states is one for each time point; and that we can reasonably model the states dynamics by a Markovian process – i.e. that the probability of a state being active depends purely on the data and which state is active in the previous time point.

Each of these patterns or states are an instantiation of a certain probability distribution. The HMM is generic in the sense that it can accommodate different state probability distributions, depending of the type of data we are processing and the features that we wish to model. A suitable state choice for fMRI data is the Gaussian distribution (Vidaurre et al., 2017), where each state, indexed by *k*, is modelled by a certain “mean activity*”* parameter *μ*_*k*_ and a covariance matrix *Σ*_*k*_. Let *x*_*t*_ be the data at time point *t* – i.e. the value of the ICA time courses at *t*. The probability density function that describes *x*_*t*_, assuming that state *k* is active at time *t*, is given by

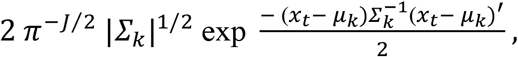

where *J* is the number of brain regions (here ICA components), |*Σ*_*k*_| is the determinant of the state-specific covariance matrix *Σ*_*k*_ and exp is the exponential function. Here, *Σ*_*k*_ represents the covariance of the residuals, i.e. after subtracting the mean parameter *μ*_*k*_ to the signal.

In this paper, each state is parametrised as Gaussian distribution with no mean parameter. Note that the interpretation of *Σ*_*k*_ as FC is not analogous to what is typically referred to when time-varying FC is assessed using sliding-window analysis (Thompson and Fransson, 2018). This is because, as opposed to sliding windows, in this type of HMM the mean *μ*_*k*_ is also allowed to be time-varying. Therefore, in order to focus the HMM decomposition on the FC changes, and in order to make the HMM estimation more comparable to standard analyses of time-varying FC, we enforced *μ*_*k*_ = 0, by describing the probability density function for state *k* as

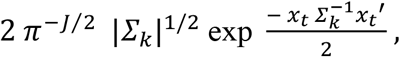

where *Σ*_*k*_ represents state-specific FC. In this model, *Σ*_*k*_ is assumed to be distributed as a Wishart distribution. Note that this is equivalent to having each state being Wishart distributed. Importantly, this model carries information of *both* time-averaged FC and time-varying FC.

Another important element of the HMM, also estimated from the data, is the transition probability matrix (TPM), which encodes the probability of transitioning from one state to another at any time point. Practically speaking, the TPM serves two purposes: it identifies the transitions that are more probable, and it regularises the state switching to minimise the amount of spurious transitions. In particular, whenever we have a more persistent (temporally regularised) solution, the diagonal elements will be comparably larger than the off-diagonal elements of the TPM.

The estimation of the HMM, carried out through a procedure of Bayesian variational inference (Wainwright and Jordan, 2008; Vidaurre et al., 2018a), was first computed at the group level, such that the state probability distributions were shared across subject – though the state time courses and the time spent in each state were subject-specific.

Next, we computed subject-specific HMMs using a process that we refer to as *dual-estimation* (in analogy to dual-regression in ICA; Beckmann et al., 2009). To do this, we simply used the subject-specific state time courses to compute a subject-specific estimation the states; then, based on these state estimations, we recomputed the state time courses and the TPM for each subject. In order to make cross-validated behavioural predictions (see below), we followed the conservative procedure of estimating the group-level HMM only on the cross-validation training sets, so that, afterwards, we could obtain the dual-estimated HMMs on both the training and testing sets.

### The HMM contains time-averaged FC information

The HMM contains information not just on time-varying FC (how FC changes within each session), but also regarding the time-averaged FC (the subject-specific FC information that remains stable across sessions for each subject). This is because it is possible to fully reconstruct the time-averaged FC estimation from the dual-estimated HMM simply by computing a weighted average of the states’ covariance (FC) matrices for each subject, where the weights are given by the fractional occupancies and the fractional occupancies are defined as the total proportion of time spent at each state for every given subject (Baker et al., 2014).

Given *N* subjects and *K* states, the group level HMM estimation represents some of the subject-specific time-averaged (or static) FC (avFC) information, according to the following expression:

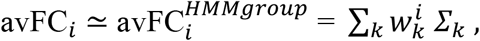

where *i* indexes subjects and 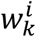 represents the fractional occupancy for subject *i* and state *k* (i.e. the total time spent on that state for that subject). Given that the number of states is lower than the number of subjects (*K* < *N*), this is an approximation, and therefore there is some time-averaged FC information that is not captured by the HMM. Likewise, the HMM has information (for example in the TPM) that is not captured by a standard time-averaged FC estimation; formally, we refer to this differential information as time-varying FC.

As opposed to the group level estimation, the dual-estimated HMM estimation captures all the time-averaged FC information

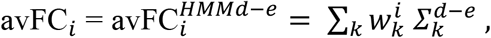

where *d-e* denotes dual-estimated. This is because:

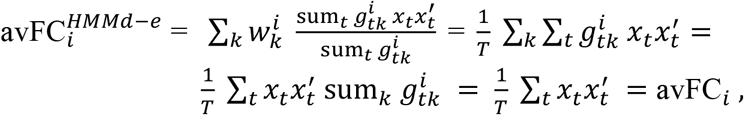

where 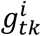 is the probability for subject *i* to be in state *k* at time point *t*, sum_*k*_ 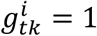 by the definition of a probability, and 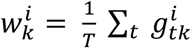.

### The HMM contains time-varying FC information

We have established that the HMM contains some time-averaged FC information. But, to which extent does the HMM capture time-varying FC information, above and beyond the time-averaged FC?

As a sanity check, in **Fig SI-5** we show that the time-averaged FC contains information that is essentially uncorrelated to the FC temporal variability. To compute a measure of the extent to which there is time-varying FC for each pair of regions, we first constructed an instantaneous estimate of FC at each time point, using a weighted sum of the dual-estimated HMM states’ FC, weighted by the assigned HMM state probabilities at that time point. We then took the variance of these instantaneous estimates of FC across time to produce a (regions-by-regions) matrix of estimated FC temporal variability for each given subject. We then compared this to the time-averaged FC, confirming that these are unrelated.

### Other HMMs with no time-varying FC information

Previously, we have shown that the dual-estimated HMMs contain all the information there is about time-averaged FC. Having *K* FC descriptions per subject instead of one, plus a TPM ruling the transitions between these states, it is apparent that the HMM contains additional information beyond time-averaged FC. An important question is then what that additional information represents. There are three possible sources of variability: actual within-session changes of FC (i.e. time-varying FC), within-session changes in the variance of the signal, and estimation noise. By meaningfully relating the HMM information to behaviour above and beyond the influence of time-averaged FC (see below, and Results) we can rule out the possibility that the HMM extra parameters are purely noise-driven. However, given that both variances and correlations (i.e. FC) are part of the state descriptions, there is no straightforward analytical way to disambiguate how much these two elements drove the inference of the HMM. In order to prove the relevance of pure time-varying FC in the HMM estimation, we obtained alternative HMM estimations where the states are purely derived by changes in the variance of the signal. The probability density function of this model is given by

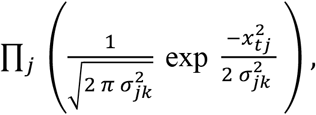

where ∏_*j*_ (·) represents multiplication across regions,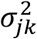 is the variance for region *j* and state *k*, and *x*_*tj*_ is the value of the signal for region *j* at time point *t*.

The fact that HMM using full covariances matrices can explain aspects of behaviour that this model was unable to explain (see **Fig 4**) suggests that there is relevant information in the HMM that is not related to changes in variance.

Even though the mean parameter of the Gaussian distribution (which reflects the amount of activity of each state with respect to the average signal) was not included in the model in the first place, we estimated a third HMM model where the states were solely defined by the mean, i.e. without state-specific covariances – and with a shared, global covariance. The purpose of this analysis is to rule out the possibility that this type of information, though not explicitly included in the HMM description used here, permeated the state covariance matrices and determined the HMM inference. The probability density function of this model is given by

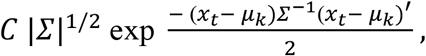

with one single covariance matrix *∑* shared across states. Note that this model holds important similarities with ICA, in the sense that each state or component is represented by a map of activation. Again, as reflected in **Fig 4**, this model is less predictive of behaviour.

### Measures of structural variability

We applied independent component analysis (implemented by the Melodic tool in FSL; Jenkinson et al., 2012) on the fractional anisotropy (FA), mean diffusivity (MD), and voxel-based morphometry (VBM) values for each subject across the whole brain (2mm resolution); resulting in 50 independent components of FA, MD, and VBM variability across subjects.

In more detail, the structural T1 weighted data was preprocessed using the computational analysis toolbox (CAT)−12 (Nenadic et al., 2015), which extends the SPM’s VBM pipeline (Ashburner and Friston, 2000). Before grey matter volume estimation, all participants’ T1 images were affinely aligned, segmented, normalized, and bias-field-corrected, yielding images containing grey and white matter segments and CSF. DARTEL (Ashburner, 2007) was then used to normalize all images to a standard grey matter template provided by CAT-12. Subsequently, all grey matter volumes were smoothed with a 9.4 mm FWHM Gaussian smoothing kernel (sigma = 4 mm). The diffusion weighted data was preprocessed using the DTIFIT routine from FSL (Jenkinson at al., 2012) in order to extract FA and MD. More details about structural preprocessing can be found in Llera et al. (2019).

### Measuring dissimilarities between subjects

The kernel-based prediction algorithm employed in this paper is based on distance matrices (DM) containing the dissimilarities between each pair of subjects within the geometrical space defined by each type of representation (see **Fig 1**). As mentioned, the main purpose of this approach here is to abstract ourselves from the specifics of each representation (e.g. time-averaged or time-varying FC) and their complexity, so that the prediction is made in a comparable fashion. Furthermore, there is not a straightforward way to unwrap the parameters of an HMM model into a vector of predictive features, so that a standard regression model can be applied. Because it is possible to compute distances between HMM models more straightforwardly, a kernel-based approach is a more natural way to make predictions in this case.

We next detail how to compute DMs in the spaces defined by the different imaging-based modalities: time-varying FC, time-averaged FC, and structural.

We first discuss the HMM model, which, as discussed above, contains information about both the time-averaged FC and time-varying FC. In particular, we computed the symmetric Kullback-Leibler divergence between each pair of (dual estimated) subject HMMs, denoted as *M*^*1*^ and *M*^*2*^.

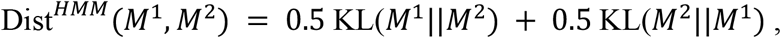

where KL(*M*^*1*^ ‖ *M*^*2*^) represents the standard (non-symmetric) Kullback-Leibler divergence between probabilistic models *M*^*1*^ and *M*^*2*^. More specifically, the Kullback-Leibler divergence represents how much information a probability distribution contains in relation to a second reference probability distribution. Whereas the Kullback-Leibler divergence has a closed form for various well-known distributions (e.g. the Gaussian distribution), this is not the case for more complex probability distributions such as the one represented by the HMM. For this reason, we adapted the mathematical approximation proposed by Do (2003) for discrete state distributions to the Gaussian case:

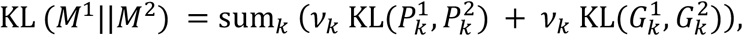

where 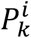 represents the (Dirichlet-distributed) probabilities of transitioning from state *k* to any of the other states according to model *i* (i.e. the *k*-th row of the TPM); 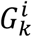 is the state Gaussian distribution for state *k* and model *i*; and *ν*_k_ is a factor representing the weight of state *k* in *M*^*1*^ (see below). Given the initial probabilities *π*^*1*^ of the HMM state time courses for model *M*^*1*^ (which are computed from the data together with the TPM), *ν* can be numerically computed such that it meets the following necessary criteria (see Do, 2003):

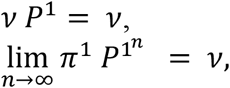

The expressions for 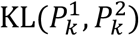 and 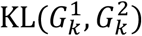 are standard and can be found elsewhere (MacKay 2003). The code to compute the symmetric Kullback-Leibler divergence between two HMM models is provided in^1^. Note that these expressions require the states to be matched between HMM models; i.e. the first state of *M*^1^ must correspond to the first state of *M*^2^. This is guaranteed here by the fact that the dual-estimated HMMs are derived from the same group-level HMM.

The second type of DM corresponds to the time-averaged FC. To keep the comparisons fair, and in line with the approach taken for the time-varying FC, we described the time-averaged FC by fitting a Gaussian distribution per subject. Given that the time series were standardised for each subject (i.e. they are demeaned and have variance equal to 1.0), the resulting Gaussian distributions only contain a covariance matrix that is mathematically equivalent to using a Pearson’s correlation matrix. The time-averaged FC’s DM was computed using the symmetric Kullback-Leibler divergence between each pair of the subject-specific Gaussian distributions,

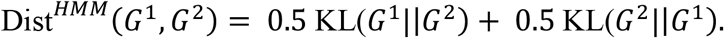

Note that, because this way we are taking into account the non-Euclidean geometry of the covariance matrices, this approach is mathematically more principled and therefore statistically more efficient than using correlations across the off-diagonal elements of the FC matrices (as is more commonly done in the literature).

Finally, the third type of DM is computed from the structural information, using the three considered structural measures: FA, MD and VBM. As discussed earlier, we have 50 ICA components for each measure, so the data consist of 50 weights per subject in each case. Given that there is no specific geometrical constrain on these values, we just used Euclidean distances between each pair of subjects in order to estimate the corresponding DMs.

### Predicting subject traits using kernel ridge regression

Our goal is to capture the relationship between the representations of the brain imaging data (time-varying FC, time-averaged FC, and structural information) and subject traits. One way to achieve this is by regressing a set of nonlinear mappings of the imaging-derived features onto the behavioural traits, e.g.:

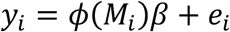

where *y*_*i*_ is the (*N* by 1) behavioural trait of subject *i*, and *ϕ*(*M*_*i*_) is a nonlinear function that operates on a representation *M*_*i*_ of the brain imaging data. To implement this regression model, we need to specify the choice of the function *ϕ*(*M*_*i*_) and also the imaging-derived features it should operate on – a not straightforward task.

An alternative and simpler approach to working with this regression model is to take advantage of the so-called kernel trick, whereby predictions of an out-of-sample subject’s behavioural trait, *ŷ*_*i*_, are made to depend on a kernel function without the need of manually defining *ϕ*(*M*_*i*_) (Schölkopf and Smola, 2001). Specifically, we use kernel ridge regression (KRR), which is formulated as

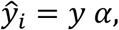

where *α* is a (*N* by 1) vector of KRR weights, and *y* represent the observed (*N* by 1) vector of values of the behavioural trait from the training CV-fold. As shown in the mathematical derivations by Saunders et al. (1998), we can make use of the kernel trick to estimate *α* as

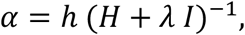

where *λ* is a regularisation parameter. As with other kernel-based approaches, such as the support vector machine or Gaussian processes, KRR works on a (*N* by *N*; where *N* is the number of subjects) kernel matrix *H*, which is computed by applying some kernel function on the corresponding DM. Here, we chose a Gaussian radial basis function kernel, parametrised by a radius parameter τ (Hastie et al., 2011):

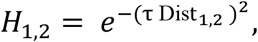

where Dist_1,2_ is the distance between the representation for two different subjects within the training CV-fold, and *H*_1,2_ is the corresponding element of the kernel matrix. That is, once we have computed the corresponding DM, the KRR approach does not need to consider where these distances come from. The choice of the Gaussian kernel function is motivated by the fact that it generalises well to most domains, given its lack of strong assumptions (Schölkopf and Smola, 2001). On these grounds, *h* is a (1 by *N*) vector containing the result of applying the Gaussian kernel to the *N* distances between each of the subjects in the training set and subject *i* in the test CV-fold.

In summary, the KRR formulation (also benchmarked in a neuroimaging context against deep learning methods by He et al., 2020) is equivalent to having a nonlinear prediction using an explicit nonlinear function *ϕ*(*M*_*i*_), but without having to directly design, use, or even know, such function; instead, we only need to specify a distance measure between the representations (e.g. of the HMMs, the structural images, or the time-averaged FC matrices) and a valid kernel function. The code for KRR, which uses a nested cross-validation loop to select both *λ* and τ, is provided in ^2^.

### Accounting for brain structure in the predictions

A central goal of this paper is to assess how the predictive power of the time-averaged and time-varying FC representations relates to the anatomy. For this purpose, we used cross-validated KRR to estimate FA-, DM- and VBM-based predictions for each behavioural trait, using their respective DMs. That is, we estimated regularised KRR coefficients on the training folds and applied them on each testing fold, in turn, to eventually produce an (*N* by 1) vector of predicted traits for each behavioural variable and structural modality. Then, we computed the corresponding residuals as the difference between the predicted and the empirical traits, and used these as FA-, MD- or VBM-deconfounded behavioural traits in the subsequent time-averaged-FC-based and HMM-based predictions. Cross-validation-based deconfounding was chosen because it is less aggressive and biased than standard deconfounding (Snoek et al., 2019).

### Motion correction

Since motion is known to influence both the estimation of time-varying FC and the prediction of behavioural variables, we used FIX and confound regressors at the level of the individual subject time series (Smith et al., 2013b). Furthermore, we included the derived motion parameters as confounds in the KRR prediction in order to also control for between-subject differences in motion.

## Acknowledgements

DV is supported by a Novonordisk Emerging Investigator Award (NNF19OC-0054895). MWW’s research is supported by the NIHR Oxford Health Biomedical Research Centre, by the Wellcome Trust (106183/Z/14/Z), and the MRC UK MEG Partnership Grant (MR/K005464/1). The Wellcome Centre for Integrative Neuroimaging is supported by core funding from the Wellcome Trust (203139/Z/16/Z). AL is supported by the Horizon2020 Programme CANDY (Grant No. 847818).

## Supplemental Information

**Fig SI-1.**
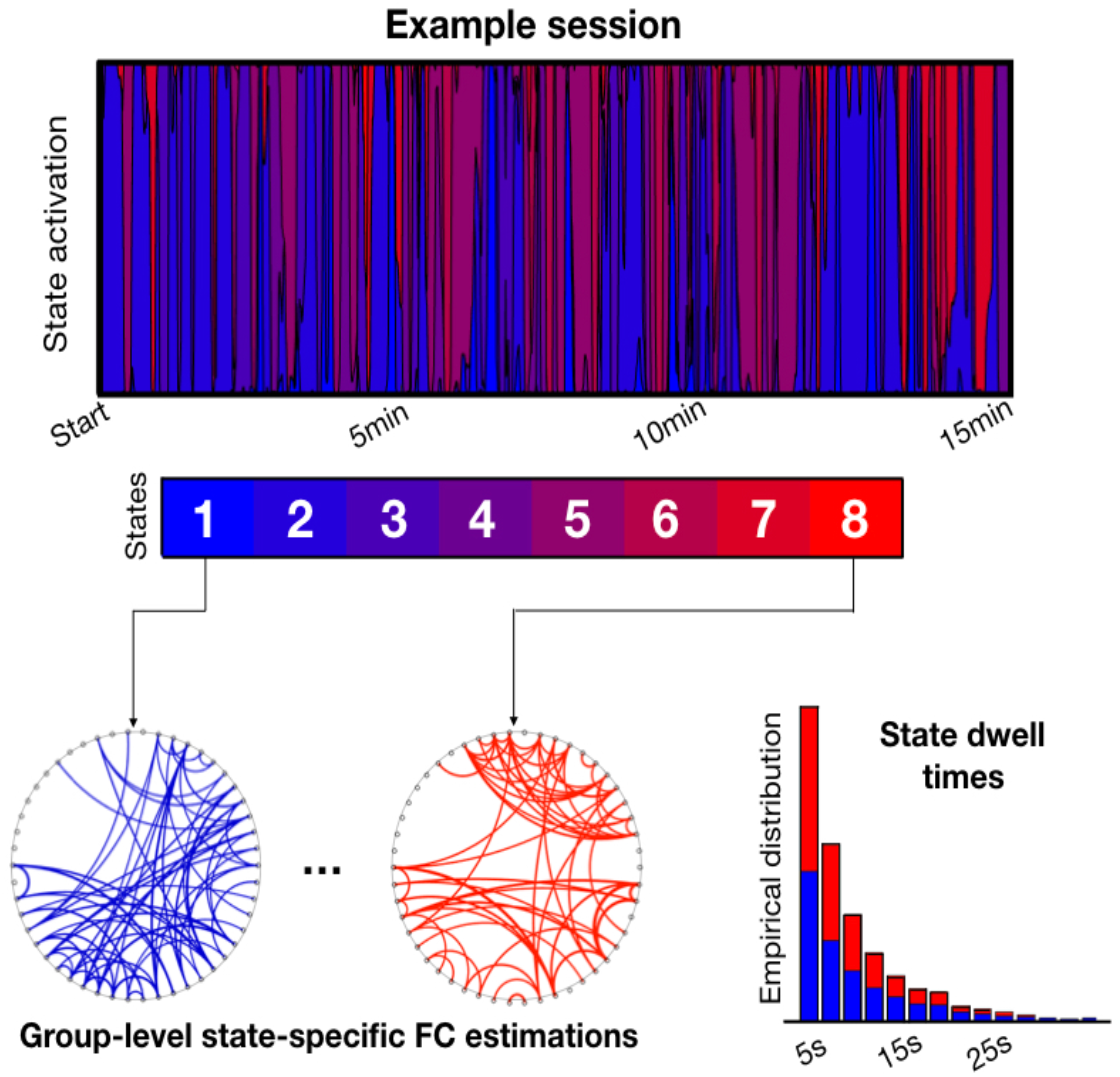
Illustration of the HMM description of time-varying FC, for an example session; states are represented as connectivity matrices. On the bottom right, depiction of the distribution of state dwell times for two of the states, at the group level.

**Fig SI-2.**
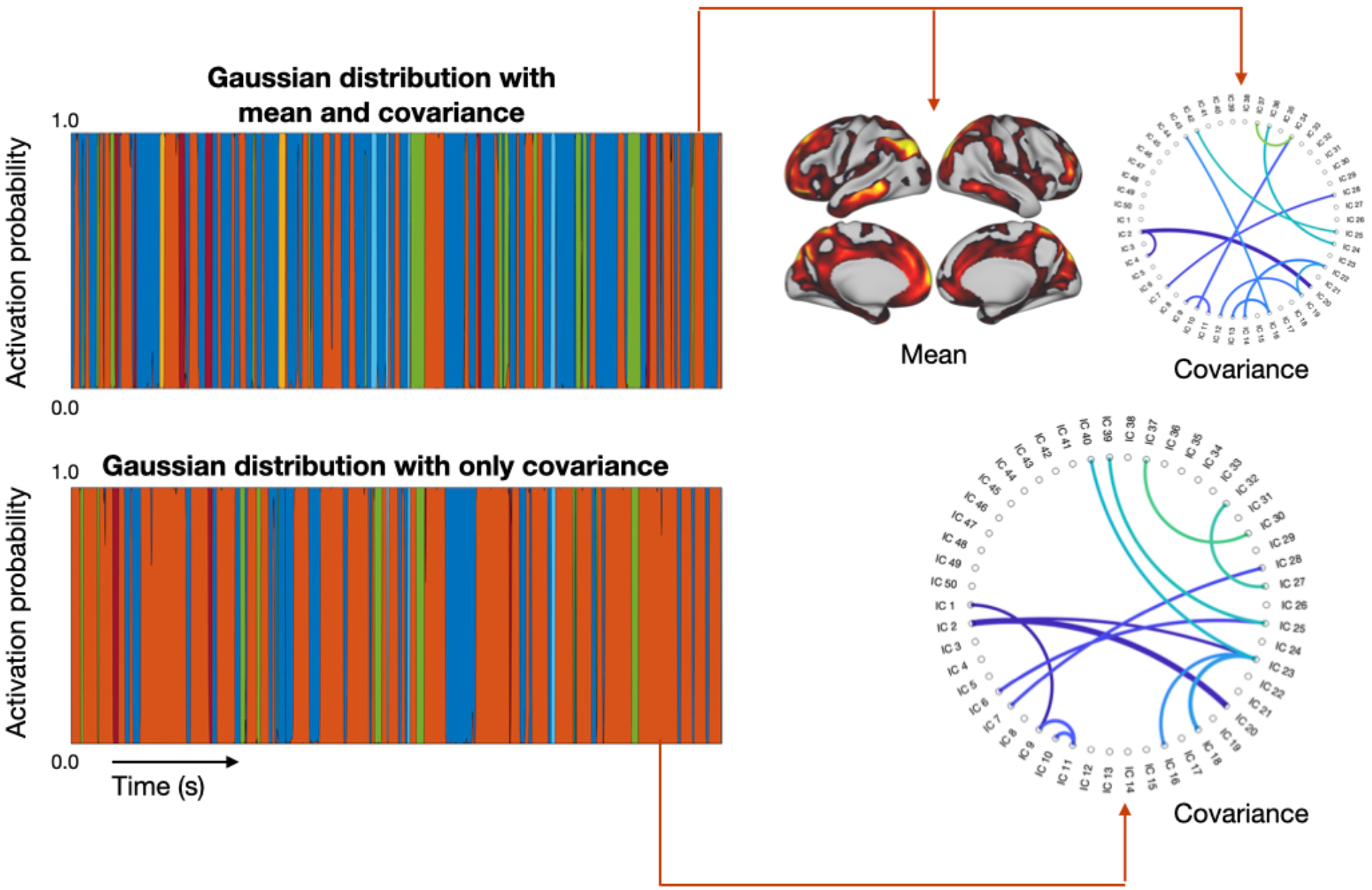
Example of the HMM run on just one subject, for two varieties of the HMM: The standard HMM with states defined by Gaussian distribution with mean and covariance (Vidaurre et al., 2017), and the FC-HMM used here which only uses covariance. On the left, state time courses for one session; on the right, observation model for one state. The random seeds were equal for both runs.

**Fig SI-3.**
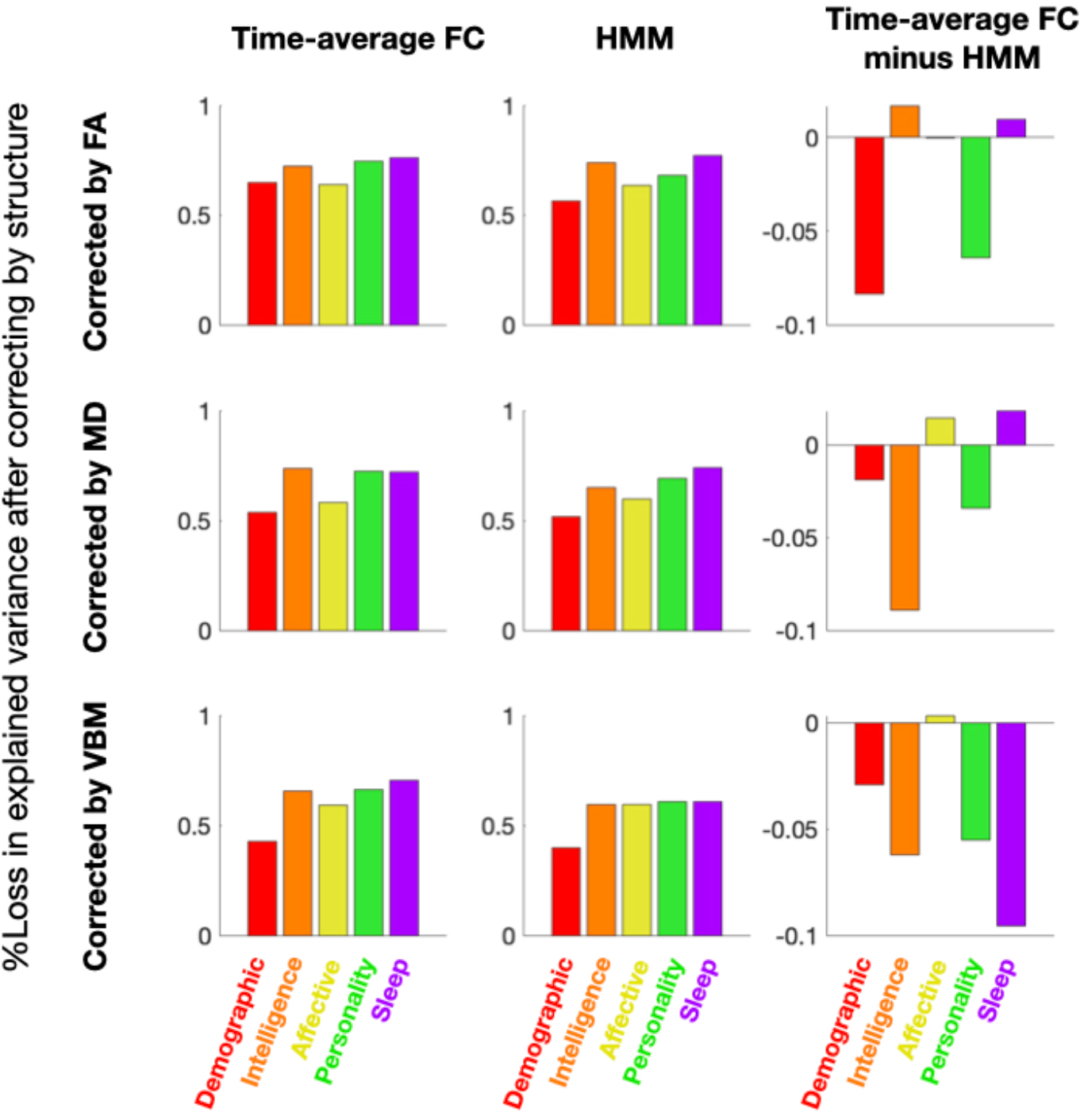
Loss in explained variance of the deconfounded predictions with respect to the non-deconfounded predictions, expressed as a percentage.

**Fig SI-4.**
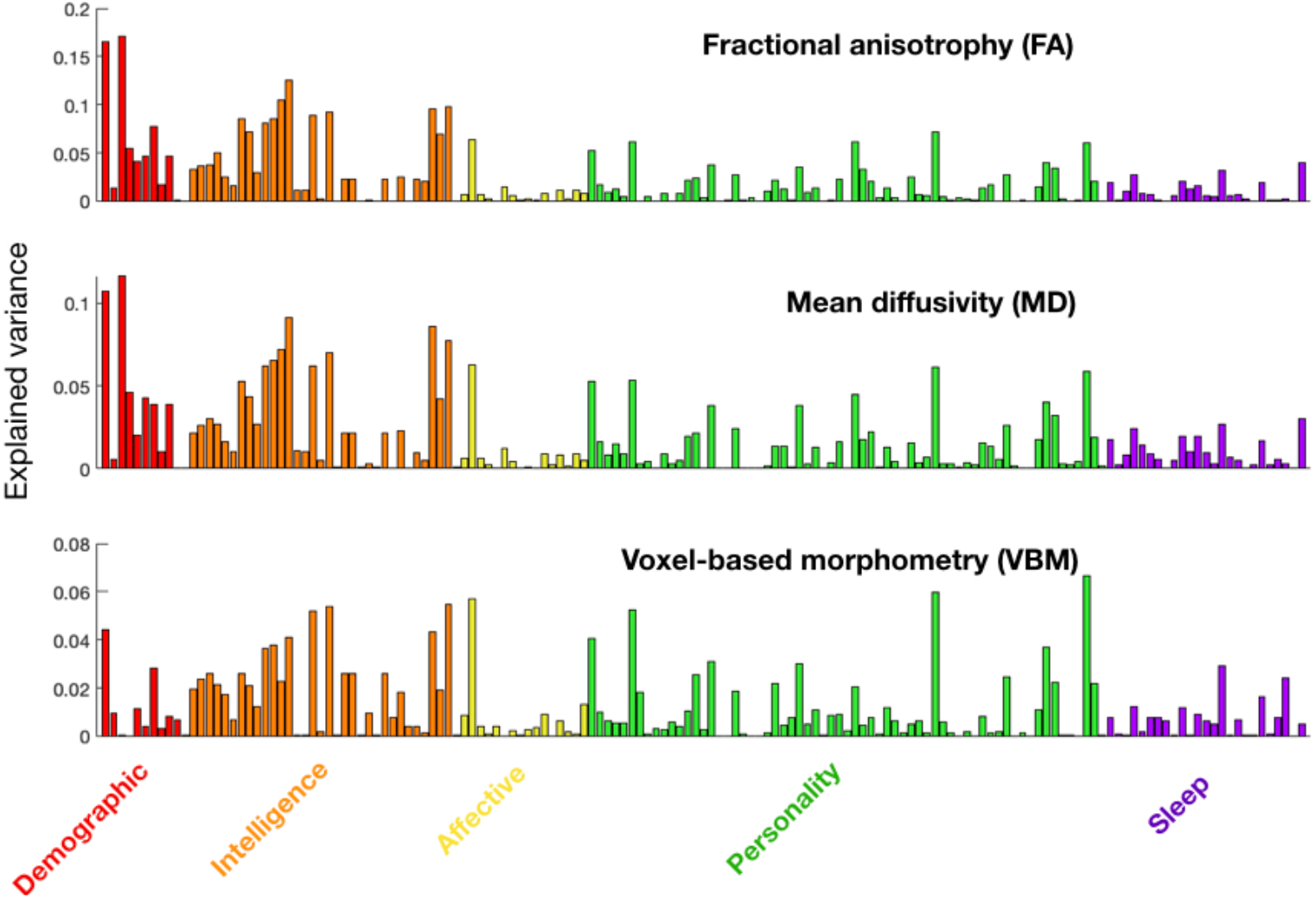
Explained variance *r*^*2*^ (in terms of squared Pearson’s correlation) of each of behavioural trait by the structural information. Traits are coloured according to five different behavioural groups: demographics, intelligence, affective, personality and sleep.

**Fig SI-5.**
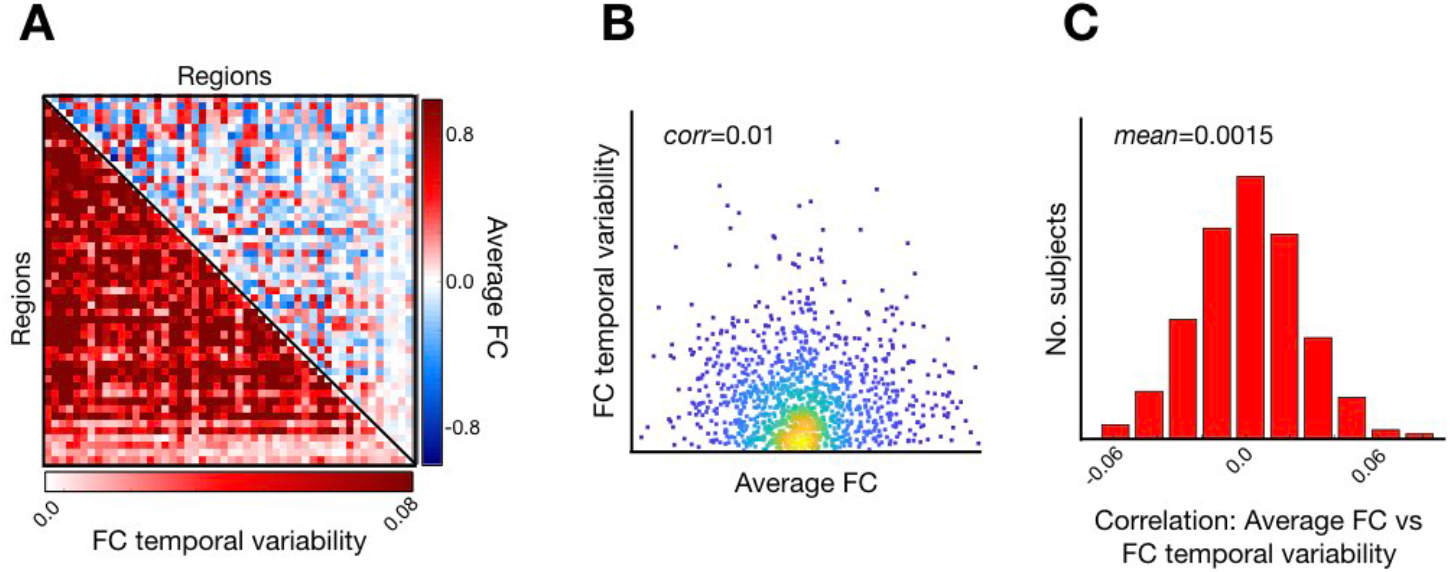
In order for FC temporal variability and time-averaged FC to explain distinct aspects of behaviour, these representations must contain non-shared elements of information of brain function. This amounts to showing that there is unique, subject-specific behaviourally-relevant information in the time-varying FC, which is not contained in the time-averaged FC. To do this, we computed the time-averaged FC for each subject and compared it with a measure of FC temporal variability for each subject (see Methods). This figure shows that the amount of time-averaged FC for any pair of regions is unrelated to amount of FC temporal variability for such a pair: (**A)** an example for one subject, where the upper triangular matrix represents time-averaged FC and the lower triangular matrix represents FC temporal variability; **(B)** the relation between time-averaged FC and FC temporal variability for that same subject as a scatter plot, where each dot corresponds to a pair of regions. Although the null hypothesis cannot be proved in this way, we note that the correlation between these measures is 0.01 and non-significant; **(C)** histogram of correlations between time-averaged FC and FC temporal variability across subjects; the mean correlation is 0.0015 and is non-significantly positive.

**Table SI-1.**
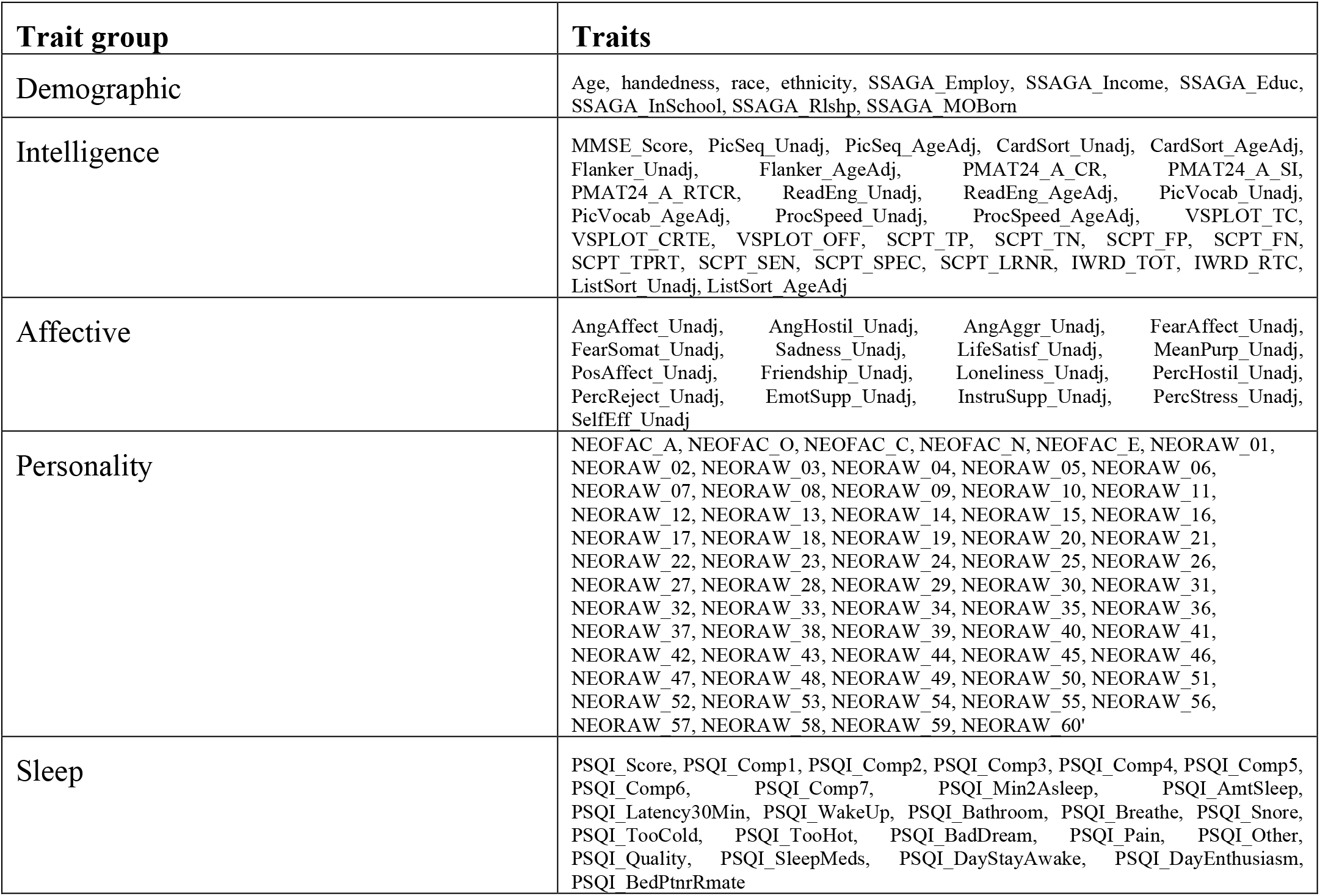
List of behavioural and anatomical traits per group.

## Footnotes

1 https://github.com/OHBA-analysis/HMM-MAR/blob/master/utils/math/hmm_kl.m

2 https://github.com/OHBA-analysis/HMM-MAR/blob/master/utils/prediction/predictPhenotype_CVHMM.m

## References

1. T.W. Allan, S.T. Francis, C. Caballero-Gaudes, P.G. Morris, E.B. Liddle, P.F. Liddle, M.J. Brookes and P.A. Gowland (2015). Functional Connectivity in MRI Is Driven by Spontaneous BOLD Events. PLoS One, e0124577.

2. J. Ashburner (2007). A fast diffeomorphic image registration algorithm. NeuroImage 38, 3031–3054.

3. A.P. Baker, M. J. Brookes, I. A. Rezek, S.M. Smith, T.E.J. Behrens, P. J. P. Smith & M. W. Woolrich (2014). Fast transient networks in spontaneous human brain activity. eLife 3: e01867.

4. C. Baldassano, J. Chen, A. Zadbood, J.W. Pillow, U. Hasson and K.A. Norman (2017). Discovering event structure in continuous narrative perception and memory. Neuron 95, 709–721.

5. P.J. Basser and C. Pierpaoli (1996). Microstructural and physiological features of tissues elucidated by quantitative-diffusion-tensor MRI. Journal of Magnetic Resonance 213, 560–570.

6. P.J. Basser, J. Mattiello, D. Le Bihan (1994). Estimation of the effective self-diffusion tensor from the NMR spin-echo. Journal of Magnetic Resonance, Series B 103, 247–254.

7. J. Ashburner and K.J. Friston (2000). Voxel-based morphometry – The methods. Neuroimage 11, 805–821.

8. C.F. Beckmann, C.E. Mackay, N. Filippini and S.M. Smith (2009). Group comparison of resting-state fMRI data using multi-subject ICA and dual regression. Neuroimage 47, 148–156.

9. J. Berkson (1946). Limitations of the Application of Fourfold Table Analysis to Hospital Data. Biometrics Bulletin 2, 47–53.

10. J.D. Bijsterbosch, M.W. Woolrich, M.F. Glasser, E.C. Robinson, C.F. Beckmann, D.C. Van Essen, S.J. Harrison and S.M. Smith (2018). The relationship between spatial configuration and functional connectivity of brain regions. eLife 7, e32992.

11. B. Biswal, F. Zerrin-Yetkin, V.M. Haughton and J.S. Hyde (1995). Functional connectivity in the motor cortex of resting human brain using echo-planar MRI. Magnetic Resonance in Medicine 34, 537–541.

12. D. Bzdok, G. Varoquaux, O. Grisel, M. Eickenberg, C. Poupon and B. Thirion (2016). Formal models of the network co-occurrence underlying mental operations. PLoS Computational Biology.

13. J.S. Damoiseaux, S.A.R.B. Rombouts, J. Barkhof, P. Scheltens, C.J. Stam, S.M. Smith and C.F. Beckmann. Consistent resting-state networks across healthy participants. Proceedings of the National Academy of Sciences of the USA 103, 13848–13853 (2006).

14. E. Duff, T. Makin, M. Cottaar, S.M. Smith and M.W. Woolrich (2018). Disambiguating brain functional connectivity. NeuroImage 173, 540–550.

15. M. N. Do (2003). Fast approximation of Kullback–Leibler distance for dependence trees and hidden Markov Models. IEEE Signal Processing Letters 10, 115–118.

16. E. Efron and R. Tibshirani (1986). Bootstrap methods for standard errors, confidence intervals, and other measures of statistical accuracy. Statistical Science 1, 54–74.

17. E.S. Finn, X. Shen, D. Scheinost, M.D. Rosenberg, J. Huang, M.M. Chun, X. Papademetris and R.T. Constable (2015). Functional connectome fingerprinting: identifying individuals using patterns of brain connectivity. Nature neuroscience 18, 1664–1671.

18. M.D. Fox and M.E. Raichle. Spontaneous fluctuations in brain activity observed with functional magnetic resonance imaging. Nature Reviews Neuroscience 8, 700–711.

19. L.L.E. Frank and J.H. Friedman (1993). A statistical view of some chemometrics regression tools. Technometrics 7, 109–135

20. C. Gratton, T.O. Laumann, A.N. Nielsen, D.J. Greene, E.M. Gordon, A.W. Gilmore, S.M. Nelson, R.C. Coalson, A.Z. Snyder, B.L. Schlaggar, N.U.F. Dosenbach and S.E. Petersen (2018). Functional brain networks are dominated by stable group and individual factors, not cognitive or daily variation. Neuron 98, 439–452.

21. A.S. Greene, S. Gao, D. Scheinost and R.T. Constable (2018). Task-induced brain state manipulation improves prediction of individual traits. Nature Communications 9, 2807.

22. L. Griffanti et al (2014). ICA-based artefact removal and accelerated fMRI acquisition for improved resting state network imaging. NeuroImage 95, 232–247.

23. M. Hampson, N.R. Driesen, P. Skudlarski, J.C. Gore, and R.T. Constable (2006). Brain connectivity related to working memory performance. The Journal of Neuroscience 26, 13338–13343.

24. S.J. Harrison, M.W. Woolrich, E.C. Robinson, M.F. Glasser, C.F. Beckmann, M. Jenkinson and S.M. Smith (2015). Large-scale probabilistic functional modes from resting state fMRI. NeuroImage 109, 217–231.

25. U. Hasson, H.C. Nusbaum and S.L. Small (2009). Task-dependent organization of brain regions active during rest. Proceedings of the National Academy of Sciences of the USA 106, 10841–10846.

26. T. Hastie, R. Tibshirani and J. Friedman (2001). The Elements of Statistical Learning. Springer, 2001.

27. T. He, R. Kong, A.J. Holmes, M. Nguyen, M.R. Sabuncu, S.B. Eickhoff, D. Bzdok, D. Feng and B.T. Yeo (2020). Deep neural networks and kernel regression achieve comparable accuracies for functional connectivity prediction of behavior and demographics. Neuroimage 206, 116276.

28. J.F. Hipp, A.K. Engel and M. Siegel (2011). Oscillatory synchronization in large-scale cortical networks predicts perception. Neuron 69, 387–396.

29. J.F. Hipp and M. Siegel (2015). BOLD fMRI correlation reflects frequency-specific neuronal correlation. Current Biology 25, 1368–1374.

30. O. Jensen, K. Kaise and J.P. Lachaux (2007). Human gamma-frequency oscillations associated with attention and memory. Trends in Cognitive Sciences 30, 317–324.

31. M. Jenkinson, C.F. Beckmann, T.E.J. Behrens, M.W. Woolrich and S.M. Smith (2012). FSL. NeuroImage 62, 782–790.

32. R. Kong, J. Li, C. Orban, M.R. Sabuncu, H. Liu, A. Schaefer, N. Sun, X.N. Zuo, A.J. Holmes, S.B. Eickhoff and B.T. Yeo (2019) Spatial topography of individual-specific cortical networks predicts human cognition, personality, and emotion. Cerebral Cortex 29, 2533–2551.

33. A. Kucyi, M.J. Hove, M. Esterman, R.M. Hutchison and E.M. Valera (2017). Dynamic brain network correlates of spontaneous fluctuations in attention. Cerebral Cortex 25, 1831–1840.

34. A. Kucyi, A. Tambini, S. Sadaghiani, S. Keilholz and J.R. Cohen (2018). Spontaneous cognitive processes and the behavioral validation of time-varying brain connectivity. Network Neuroscience 2, 397–417.

35. A. Kucyi (2017). Just a thought: How mind-wandering is represented in dynamic brain connectivity. NeuroImage 180, 505–514.

36. R. Liégeois, T.O. Laumann, A.Z. Snyder, J. Zhou and B.T. Yeo (2017) Interpreting temporal fluctuations in resting-state functional connectivity MRI. NeuroImage 163, 437–455.

37. R. Liégeois, J. Li, R. Kong, C. Orban, D. Van De Ville, T. Ge, M.R. Sabuncu and B.T. Yeo (2019). Resting brain dynamics at different timescales capture distinct aspects of human behavior. Nature Communications 10, 2317.

38. A. Llera, T. Wolfers, P. Mulders and C.F. Beckmann (2019). Inter-individual differences in human brain structure and morphometry link to variation in demographics and behavior. eLife 8, e44443.

39. D. Lurie, D. Kessler, … and V. Calhoun (2020). Questions and controversies in the study of time-varying functional connectivity in resting fMRI. Network Neuroscience 4, 30–69.

40. D.J.C. MacKay (2003). Information Theory, Inference, and Learning Algorithms. Cambridge University Press.

41. S. Palva and J.M. Palva (2012). Discovering oscillatory interaction networks with M/EEG: challenges and breakthroughs. Trends in Cognitive Sciences 16, 219–230.

42. U. Pervaiz, D. Vidaurre, M.W. Woolrich and S.M. Smith (2020). Optimising network modelling methods for fMRI. NeuroImage 211, 116604.

43. G.C. O’Neill, P.K. Tewarie, G.L. Colclough, L.E. Gascoyne, B.A.E. Hunt, P.G. Morris, M.W. Woolrich, M.J. Brookes (2017). Measurement of dynamic task related functional networks using MEG. NeuroImage 146, 667–678.

44. I. Nenadic, R. Maitra, K. Langbein, M. Dietzek, C. Lorenz, S. Smesny, J.R. Reichenbach, H. Sauer and C. Gaser (2015). Brain structure in schizophrenia vs. psychotic bipolar I disorder: a VBM study. Schizophrenia Research 165, 212–219.

45. L. Nickerson, S.M. Smith, D. Öngür and C.F. Beckmann (2017). Using Dual Regression to Investigate Network Shape and Amplitude in Functional Connectivity Analyses. Frontiers in Neuroscience 11, 115.

46. A.J. Quinn, D. Vidaurre, R. Abeysuriya, R. Becker, A.C. Nobre and M.W. Woolrich (2018). Task-Evoked Dynamic Network Analysis Through hidden Markov modeling. Frontiers in Neuroscience 12, 603.

47. L.R. Rabiner (1989). A tutorial on hidden Markov models and selected applications in speech recognition. Proceedings of the IEEE 77, 257–286.

48. S. Sadaghiani, J.B. Poline, A. Kleinschmidt, and M. D’Esposito (2015)). Ongoing dynamics in large-scale functional connectivity predict perception. Proceedings of the National Academy of Sciences of the USA 112, 8463–8468.

49. C. Saunders, A. Gammerman and V. Vovk (1998). Ridge regression learning algorithm in dual variables. In Proceedings of the 15th International Conference on Machine Learning.

50. B. Schölkopf and A.J. Smola (2001). Learning with Kernels. MIT Press.

51. H. Shappell, B.S. Caffo, J.J. Pekar and M.A. Lindquist (2019). Improved state change estimation in dynamic functional connectivity using hidden semi-Markov models. NeuroImage 191, 243–257.

52. B. Schölkopf and A.J. Smola (2001). Learning with Kernels. MIT Press.

53. J. Smallwood and J.W. Schooler (2015). The science of mind wandering: empirically navigating the stream of consciousness. Annual Review of Psychology 66, 487–518.

54. S.M. Smith, C.F. Beckmann, J. Andersson, E.J. Auerbach, J. Bijsterbosch, G. Douaud, E. Duff, D.A. Feinberg, L. Griffanti, M.P. Harms, M. Kelly, T. Laumann, K.L. Miller, S. Moeller, S. Petersen, J. Power, G. Salimi-Khorshidi, A.Z. Snyder, A.T. Vu, M.W. Woolrich, J. Xu, E. Yacoub, K. Ugurbil, D.C. Van Essen and M.F. Glasser (2013b). Resting-state fMRI in the Human Connectome Project. Neuroimage 80, 144–168.

55. S.M. Smith, T.E. Nichols, D. Vidaurre, A.M. Winkler, T.E.J. Behrens, M.F. Glasser, K. Ugurbil, D.M. Barch, D.C. Van Essen and K.L. Miller (2015). A positive-negative mode of population covariation links brain connectivity, demographics and behaviour. Nature Neuroscience 18, 1565–1567.

56. S.M. Smith, D. Vidaurre, M. Glasser, A. Winkler, P. McCarthy, E. Robinson, X. Chen, W. Horton, M. Jenkinson, E. Duff, C. Beckmann, M.W. Woolrich, D. Marcus, D. Barch, K. Ugurbil, T. Nichols and D. Van Essen (2016). Second beta-release of the HCP Functional Connectivity MegaTrawl. https://db.humanconnectome.org/megatrawl/HCP820_MegaTrawl_April2016.pdf.

57. S.M. Smith, D. Vidaurre, C.F. Beckmann, M.F. Glasser, M. Jenkinson, K.L. Milller, T.E. Nichols, E.C. Robinson, G. Salimi-Khorshidi, M.W. Woolrich, D.M. Barch, K. Ugurbil and D.C. Van Essen (2013). Functional connectomics from resting-state fMRI. Trends in Cognitive Sciences 17, 666–682.

58. L. Snoek, S. Miletić and S. Scholte (2019). How to control for confounds in decoding analyses of neuroimaging data. NeuroImage 184, 741–760.

59. A. Stevner, D. Vidaurre, J. Cabral, K. Rapuano, S.F.V. Nielsen, E. Tagliazucchi, H. Laufs, P. Vuust, G. Deco, M.W. Woolrich, E. Van Someren and M.L. Kringelbach (2019). Discovery of key whole-brain transitions and dynamics during human wakefulness and non-REM sleep. Nature Communications 10, 1035.

60. W.H. Thompson and P. Fransson (2018). A common framework for the problem of deriving estimates of dynamic functional brain connectivity. NeuroImage 172, 896–902.

61. G.J. Thompson, W.J. Pan, M.E. Magnuson, D. Jaeger and S.D. Keilholz (2014). Quasi-periodic patterns (QPP): Large-scale dynamics in resting state fMRI that correlate with local infraslow electrical activity. NeuroImage 84, 1018–1031.

62. D. Vidaurre, A.J. Quinn, A.P. Baker, D. Dupret, A. Tejero-Cantero and M.W. Woolrich (2016). Spectrally resolved fast transient brain states in electrophysiological data. NeuroImage 126, 81–95.

63. D. Vidaurre, S.M. Smith and M.W. Woolrich (2017). Brain networks are hierarchically organised in time. Proceedings of the National Academy of Sciences of the USA 114, 12827–12832.

64. D. Vidaurre, R. Abeysuriya, R. Becker, A.J. Quinn, F. Alfaro-Almagro, S.M. Smith and M.W. Woolrich (2018). Discovering dynamic brain networks from Big Data in rest and task. NeuroImage 180, 646–656.

65. D. Vidaurre, L.T. Hunt, A.J. Quinn, B.A.E. Hunt, M.J. Brookes, A.C. Nobre and M.W. Woolrich (2018b). Spontaneous cortical activity transiently organises into frequency specific phase-coupling networks. Nature Communications 9, 2987.

66. D. Vidaurre, M.W. Woolrich, A.M. Winkler, T. Karapanagiotidis, J Smallwood and T.E. Nichols (2019). Stable between-subject statistical inference from unstable within-subject functional connectivity estimates. Human Brain Mapping 40, 1234–1243.

67. M.J. Wainwright & M.I. Jordan (2008). Graphical models, exponential families and variational inference. Foundations and Trends in Machine Learning 1, 1–305.

68. A.B. Waites, A. Stanislavsky, D.F. Abbott and G.D. Jackson (2005). Effect of prior cognitive state on resting state networks measured with functional connectivity. Human Brain Mapping 24, 59–68.

69. A. Winkler, M.A. Webster, D. Vidaurre, T.E. Nichols & S.M. Smith (2015). Multi-level block permutation. NeuroImage 123, 253–268.

70. G. Zhang, B. Cai, A. Zhang, J.M. Stephen, T.W. Wilson, V.D. Calhoun and Y.-P. Wang (2019). Estimating dynamic functional brain connectivity with a sparse hidden Markov model. IEEE Transactions on Medical Imaging 39, 488–498

